# A conserved isoleucine in the binding pocket of RIG-I controls immune tolerance to mitochondrial RNA

**DOI:** 10.1101/2022.08.02.502180

**Authors:** Ann Kristin de Regt, Kanchan Anand, Katrin Ciupka, Karl Gatterdam, Bastian Putschli, David Fusshöller, Daniel Hilbig, Alexander Kirchhoff, Charlotte Hunkler, Steven Wolter, Agathe Grünewald, Christine Schuberth-Wagner, Janos Ludwig, Katrin Paeschke, Eva Bartok, Thomas Zillinger, Gregor Hagelueken, Gunther Hartmann, Matthias Geyer, Martin Schlee

**Affiliations:** Department of Clinical Chemistry and Clinical Pharmacology, University Hospital Bonn, Bonn, Germany; Institute of Structural Biology, University Hospital Bonn, Bonn, Germany; Department of Oncology, Hematology and Rheumatology, University Hospital Bonn, Bonn, Germany; Institute of Experimental Haematology and Transfusion Medicine, University Hospital Bonn, Bonn, Germany; Unit of Experimental Immunology, Department of Biomedical Sciences, Institute of Tropical Medicine, Antwerp, Belgium

## Abstract

RIG-I is a cytosolic receptor of viral RNA essential for the immune response to numerous RNA viruses. Accordingly, RIG-I must sensitively detect viral RNA yet tolerate abundant self-RNA species. The basic binding cleft and an aromatic amino acid of the RIG-I C-terminal domain(CTD) mediate high-affinity recognition of 5’triphosphorylated and 5’base-paired RNA(dsRNA). Here, we found that, while 5’unmodified hydroxyl(OH)-dsRNA demonstrated residual activation potential, 5’-monophosphate(5’p)-termini, present on most cellular RNAs, prevented RIG-I activation. Determination of CTD/dsRNA co-crystal structures and mutant activation studies revealed that the evolutionarily conserved I875 within the CTD sterically inhibits 5’p-dsRNA binding. RIG-I(I875A) was activated by both synthetic 5’p-dsRNA and endogenous long dsRNA within the polyA-rich fraction of total cellular RNA. RIG-I(I875A) specifically interacted with a long, highly structured, polyA-bearing, non-coding mitochondrial(mt) RNA, and depletion of mtRNA from total RNA abolished its activation. Altogether, our study demonstrates that avoidance of 5’p-RNA recognition is crucial to preventing mtRNA-triggered RIG-I-mediated autoinflammation.

## INTRODUCTION

The cytosolic nucleic acid receptors of the innate immune system are crucial for the initiation and control of the antiviral immune response in infected organisms. Receptor activation induces antiviral signaling pathways leading to the expression of type-I IFN, cytokines, chemokines, and antiviral effector proteins. Since endogenous nucleic acids are obviously also present in the host, pathogen-derived nucleic acids are generally sensed via their aberrant localization, e.g. cytosolic DNA, and/or the presence of uncommon/non-self structures or modifications, known as recognition motifs (Bartok and Hartmann, 2020). Effective cytoplasmic RNA recognition is particularly challenging due to the physiological presence of a variety of endogenous RNA species in the cytoplasm. Here, the need to sensitively detect pathogenic RNA has to be balanced with the potential danger of “mistakenly” sensing endogenous RNA (Ahmad et al., 2018). Whereas excessive tolerance favors the spread of infection, non-specific immune recognition of endogenous nucleic acids can lead to devastating autoinflammatory disease, such as Aicardi–Goutières Syndrome (AGS), Singleton–Merten Syndrome (SMS) and other type-I interferonopathies (Rehwinkel and Gack, 2020; Rodero and Crow, 2016).

The members of the RIG-I like receptor family (RLR) – including RIG-I itself, MDA5, and LGP2 – are capable of recognizing only a few copies of pathogenic RNA in the cytosol among an excess of endogenous RNA species, such as rRNA, tRNA, and mRNA (Schlee and Hartmann, 2016). rRNA and tRNA are highly structured and have 5’-monophosphate (5’p) termini. mRNA from higher eukaryotes bear a 5’-cap1 structure, consisting of a 5’ to 5’ triphosphate(ppp) linkage between N7-methyl-guanosine (m7Gppp), which has a 2’O-methylated penultimate nucleotide(N1) (m7GpppN(Ome), cap1). Although long RNAs with secondary structures activate MDA5 in a 5’independent manner (Schlee and Hartmann, 2016) and are also reported to activate RIG-I (Binder et al., 2011), activation of RIG-I by short dsRNA critically depends on the structure and modifications of the 5’-terminus. Blunt-ended dsRNA bearing a 5’-triphosphate (5’ppp, including cap0) or 5’-diphosphate (5’pp) is sensitively detected by RIG-I, while a cap1 structure abrogates activation (Devarkar et al., 2016; Goubau et al., 2014; Hornung et al., 2005; Schlee et al., 2009; Schuberth-Wagner et al., 2015). Moreover, 5’OH-dsRNA also activates RIG-I, albeit less potently and only at high concentrations and with slower activation kinetics (Binder et al., 2011; Marques et al., 2006). Since free 5’ppp, 5’pp, cap0, and 5’OH RNA ends are normally absent from the cytosol of non-infected vertebrate cells, they represent logical danger signals for RIG-I to detect (Bartok and Hartmann, 2020). Conversely, active avoidance of RIG-I activation by self-structures (cap1 and 5’p) is also a plausible immune discriminatory strategy. Indeed, we previously identified such a mechanism for cap1 tolerance located within the CTD of RIG-I (Schuberth-Wagner et al., 2015). However, the effect of 5’p on RIG-I activation has been controversial in the literature: While one group reported an enhancing effect for 5’p on RIG-I activation due to RNA stabilization (Takahasi et al., 2008), another, more recent, study observed that 5’p inhibited RIG-I binding and ATPase activity (Ren et al., 2019).

In the currently accepted model, prior to RIG-I activation, an inhibitory helicase subdomain (Hel-2i) binds the caspase activation and recruitment domains (CARDs), thus suppressing CARD signaling (Suppl. Fig. 1A) (Kolakofsky et al., 2012; Kowalinski et al., 2011). However, when pathogenic dsRNA binds to the flexibly connected CTD of RIG-I, the RNA ligand is oriented to form interactions with the helicase domains Hel-1 and Hel-2i, resulting in CARD displacement, homotypic CARD-CARD interaction with the adaptor protein MAVS and downstream signaling to type I IFN induction (Suppl. Fig. 1A). Based on this model, effective interaction of dsRNA with the CTD is a prerequisite of RIG-I activation. Indeed, several studies from our group and others have characterized how structural features of the CTD critically determine which RNA molecules can bind and activate RIG-I (Lu et al., 2010, 2011; Wang et al., 2010): 5’ppp/pp is recognized by a basic binding cleft (H847, K858, K861, K888) within the CTD, and the presence of a 5’-basepair enables a stacking interaction with F853 (Suppl. Fig. 1B). In the current study, we investigate the mechanism by which the 5’pRNA interaction with RIG-I is inhibited, thus preventing recognition of self RNA. We demonstrate that, while the RIG-I CTD can bind 5’ppp-dsRNA with high affinity and 5’OH-dsRNA with moderate affinity, 5’p-dsRNA-CTD binding was barely detectable. The molecular basis for this discrimination is an isoleucine residue (I875) in the RIG-I CTD that precludes 5’p-dsRNA binding (Suppl. Fig. 1B). Co-crystal structures of 5’OH-dsRNA and 5’p-dsRNA with wt and I875A-mutated RIG-I CTD, respectively, show the steric conflict of 5’p-dsRNA with I875 compared to 5’OH-dsRNA. Correspondingly, the I875A mutation completely restored RIG-I activation by 5’p-dsRNA to the level of 5’OH-dsRNA, and long-term expression of RIG-I(I875A) resulted in robust RIG-I activation by endogenous RNA. Moreover, within total RNA, we could identify mitochondrial RNA as the most active RNA species to stimulate RIG-I(I875A). Our findings indicate that this active avoidance of recognition of endogenous 5’p-RNA is crucial to preventing RIG-I mediated autoinflammation, thus characterizing a new mechanism by which RIG-I discriminates between self and non-self RNA.

## RESULTS

### 5’p blocks RIG-I-dependent immune stimulation by short and long dsRNA

Whether 5’p-dsRNA has the ability to stimulate RIG-I activity has remained controversial (Ren et al., 2019; Takahasi et al., 2008). Although Ren et al. detected weaker binding by 5’p-dsRNA than by 5’OH-dsRNA, the study did not demonstrate a significant difference in type-I-IFN induction by 5’p-dsRNA and 5’OH-dsRNA. We compared the activity of synthetic, blunt-ended 24mer and 40mer dsRNA with 5’ppp, 5’p, or 5’OH modifications at both ends. Here, using RIG-I/MDA5 deficient (RM^-/-^) T-REx-293 HEK cells expressing RIG-I(WT) and peripheral blood mononuclear cells (PBMCs) which were treated with chloroquine to block TLR7/8 activity, we found a substantial reduction of RIG-I stimulation by 5’p modification in comparison to 5’OH (Fig. 1A-C). We also tested the sequences used by (Takahasi et al., 2008), which possessed 5’p on one strand of the dsRNA, and we could not observe a significant stimulatory effect of 5’p over 5’OH-dsRNA in our cells (Suppl. Fig. 1C, 6+4). Of note, the dsRNA molecule used in the Takahasi et al has both a 5’p <u>and</u> a 5’OH terminus. In contrast, the presence of a 5’p modification at *both* ends of the dsRNA molecule almost completely abolished RIG-I activation (Suppl. Fig. 1C, “6+7”). Taken together, this indicates that the presence of one non-phosphorylated end is sufficient to activate RIG-I, while the other 5’p end might have a different, auxiliary effect, such as increasing RNA stability (as discussed by (Takahasi et al., 2008)), leading to an enhancement of RIG-I stimulation of this unique sequence in some cell types or species. To investigate the impact of 5’p modification on longer dsRNA, we compared 60mer, 100mer, and 200mer dsRNA (*in vitro* transcribed (IVT)RNA) with 5’ppp, and 5’p modifications (Fig. 1D). 5’p-dsRNA was generated by treatment of IVT RNA with polyphosphatase, a pyrophosphatase that leaves 5’p ends from 5’ppp or 5’pp. Polyphosphatase treatment completely abolished RIG-I activation for all lengths of dsRNA. However, enzymatic removal of 5’p with alkaline phosphatase (FastAP) restored immune activation, demonstrating that the loss of RIG-I activation with 5’p-dsRNA is indeed due to the 5’p terminus (Fig. 1E).

**Figure 1:**
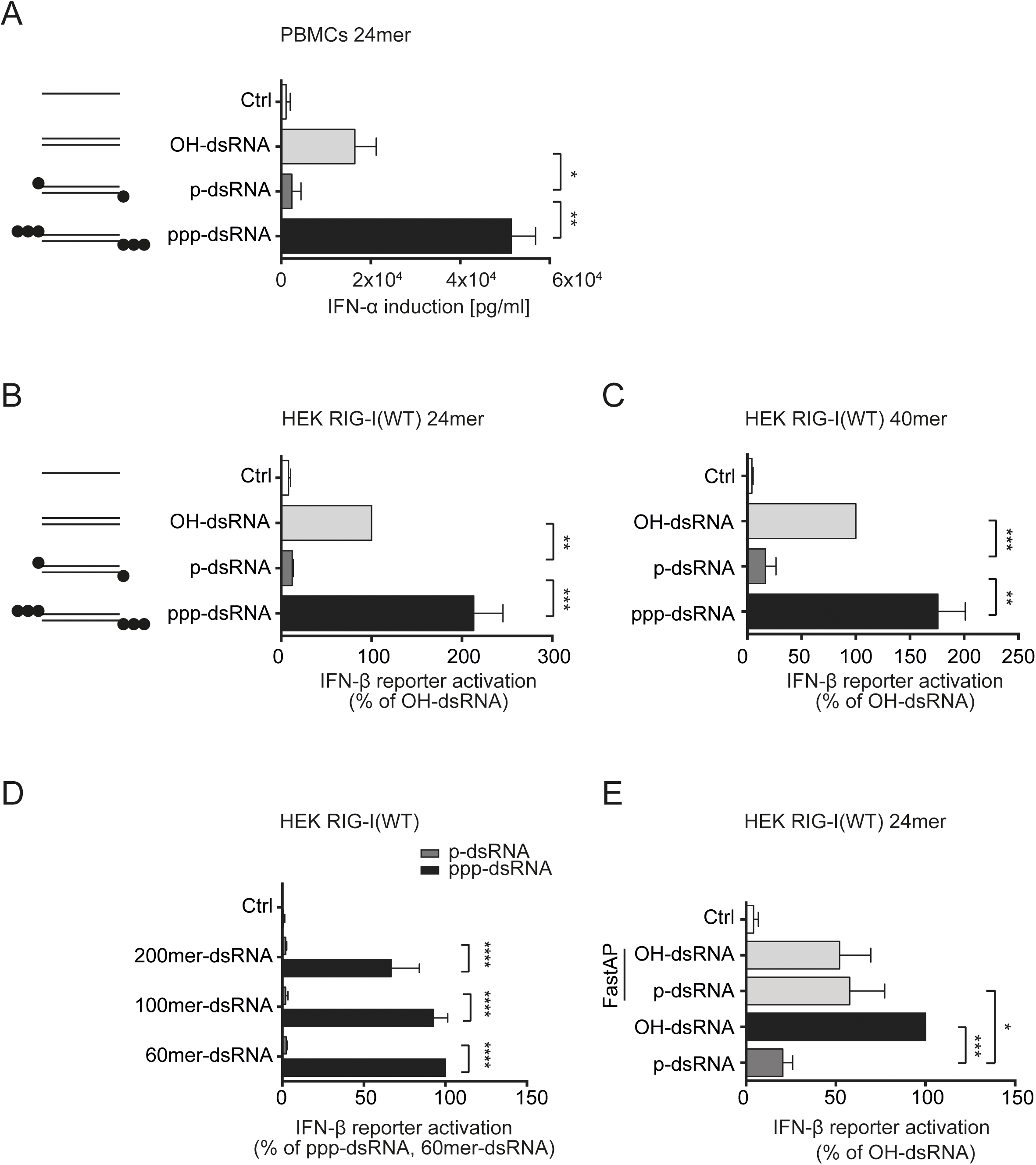
5’p blocks RIG-I-dependent immune stimulation by short and long dsRNA. A) Chloroquine pre-treated PBMCs were stimulated with the indicated synthetic RNA ligands (0.1 µg/ml). Concentrations of IFN-α in the supernatant were determined 20 h after transfection by ELISA. B,C) HEK T-REx *RIG-I*/*MDA5*(*R/M*)*^-/-^*cells expressing RIG-I(WT) were stimulated with the indicated synthetic RNA ligands (0.1 µg/ml). IFN-β reporter activation was determined 20 h after RNA transfection and is depicted as percentage of the signal of RIG-I(WT) stimulated by OH-dsRNA. A, B) 24mer dsRNAs and C) 40mer dsRNAs. D) HEK T-REx *RM^-/-^* cells expressing RIG-I(WT) were stimulated with either untreated (ppp-dsRNA) or polyphosphatase-treated (p-dsRNA) IVT-transcribed dsRNA ligands (1 ng/ml) of the indicated lengths (60, 100, or 200mer). E) HEK T-REx *RM^-/-^*cells expressing RIG-I(WT) were stimulated with the 24mer synthetic RNA ligands indicated (0.1 µg/ml). IFN-β reporter activation was determined 20 h after RNA transfection and is depicted as percentage of the signal of RIG-I(WT) stimulated with E) 60mer ppp-dsRNA. “Fast AP” indicates alkaline phosphatase treatment of the RNA prior to transfection. A-C, E) n = 4 + SD, one-way ANOVA, Dunnett’s post-test, D) n = 4 + SD, two-way ANOVA, Sidak’s post-test. Statistical significance is indicated as follows: * (p < 0.05), ** (p < 0.01), *** (p < 0.001) **** (p < 0.0001). 5’-modifications: 5’-hydroxyl (OH-), 5’-monophsphate (p-, ●), 5’-triphosphate (ppp-, ●●●). RNA sequences: Table S1.

### Isoleucine 875 in the RIG-I CTD prevents 5’p-dsRNA binding by steric exclusion

Compared to 5’OH dsRNA, 5’p-dsRNA has two additional negative charges that could theoretically facilitate its interaction with the positively charged binding cleft. Thus, it was surprising that 5’p-dsRNA was less stimulatory than dsRNA without 5’phosphate. To identify the underlying structural mechanisms of this phenomenon, amino acids in the RIG-I CTD closer than 4 Å to the α-phosphate of the RNA were identified using the crystal structure of the RIG-I CTD with 5’pp-dsRNA (3NCU;(Wang et al., 2010)). With the exception of I875, all of the other amino acids identified (namely, K858, K861, K888, and H847) have been previously shown to be involved in binding to the triphosphate, and mutation of any of these four amino acids decreases or completely abrogates immune stimulation with 5’ppp-dsRNA (Wang et al., 2010). Although, I875 is highly conserved among species (Suppl. Fig. 2), it has not been implicated in 5’ppp binding so far, and the hydrophobic side chain of isoleucine might instead have repulsive effects on charged molecules like 5’phosphates. To test this hypothesis, RIG-I/MDA5 deficient (*RM^-/-^*) HEK cells expressing either wild-type RIG-I (WT) or RIG-I with I875 mutated to alanine (I875A) from a doxycycline-inducible locus (T-REx) were stimulated with 5’p-dsRNA and 5’OH-dsRNA. Indeed, the RIG-I(I875A) mutant could be activated by 5’p-dsRNA at a level similar to 5’OH-dsRNA for both sequences tested (24mer and 40mer, Fig. 2A+B).

**Figure 2:**
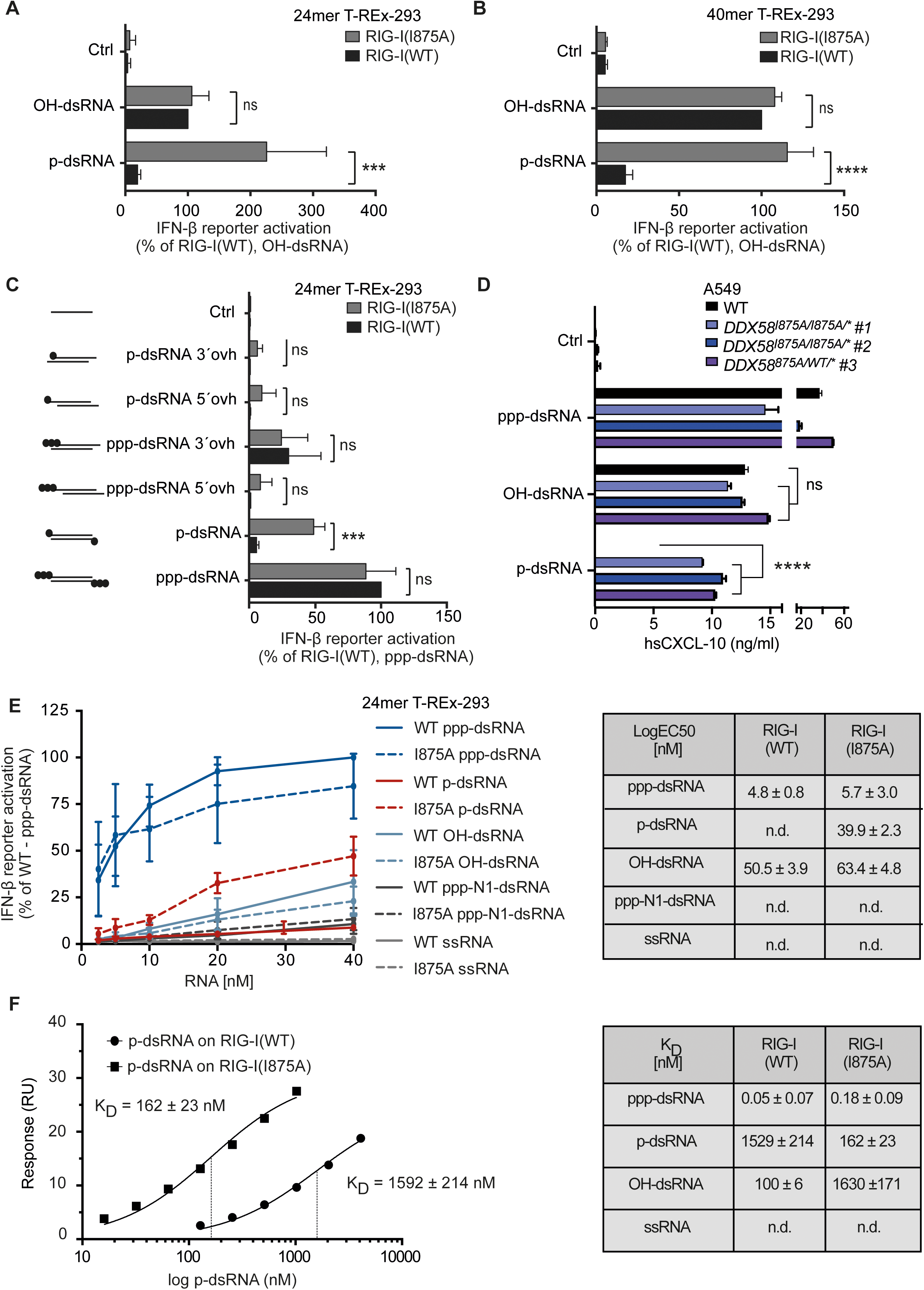
I875 in the RIG-I CTD prevents p-dsRNA binding by steric exclusion. A-C: HEK T-REx RM^-/-^ cells expressing either RIG-I(WT) or RIG-I(I875A) were stimulated with indicated synthetic RNA ligands (0.1 µg/ml). A,C) 24mer dsRNA B) 40mer dsRNA with indicated end-modifications and overhangs. IFN-β reporter activation was determined 20 h after RNA transfection and is depicted as percentage of the signal of RIG-I(WT) stimulated by A,B) OH-dsRNA or C) ppp-dsRNA. n = 3 +SD, two-way ANOVA, Sidak’s post-test. D) A549 DDX58(WT) and DDX58(I875A) gene-edited cell lines were stimulated with the RNA ligands indicated. CXCL-10 from supernatants was measured after 24 hrs. ****p<0.0001, two-way ANOVA, Dunnett’s multiple comparisons test, brackets depict 3 comparisons vs WT condition. Shown are means of duplicates + SEM of one experiment, representative of 2 independent experiments. E) RNA stimulation titration curves of HEK T-REx RM^-/-^ cells expressing RIG-I(WT) or RIG-I(I875A) stimulated with indicated RNAs (c[RNA]= 0.0625 – 1 µg/ml). IFN-β reporter activation was determined 20 h after RNA transfection and is depicted as percentage of the signal of RIG-I(WT) stimulated by ppp-dsRNA. EC50 values were calculated by nonlinear regression. n.d. = not determined (n=3 + SE). F) Dissociation constants of indicated RIG-I CTD/RNA complexes determined by surface plasmon resonance (SPR). Statistical significance is indicated as follows: *** (p < 0.001), **** (p < 0.0001), ns: not significant. 5’modifications: 5’hydroxyl (OH-), 5’monophsphate (p-), 5’triphosphate (3p-). N1-2’O-methylated 5’terminal nucleotide (N1). RNA sequences: Table S1.

To test if mutation of I875 caused a general increase in immune activation, cells were stimulated with RNA with other inhibitory features, such as 5’ or 3’ overhangs (Fig. 2C). Moreover, to exclude artefacts from overexpressed RIG-I, we then generated A549 cells with the I875A mutation in the endogenous DDX58 loci (Fig. 2D), demonstrating that RIG-I(I875) mutant protein expressed from the endogenous locus was also stimulated by p-dsRNA.

All other RNAs tested showed a similar immunostimulatory potential for wild-type RIG-I(WT) and RIG-I(I875A) cells, demonstrating the specific effect of RIG-I(I875A) 5’p recognition. Moreover, while the EC50 value for 5’p-dsRNA binding to RIG-I(WT) could not be determined owing to a lack of immune activation, the EC50 value for RIG-I(I875A) with 5’p-dsRNA was comparable to that of 5’OH-dsRNA in RIG-I(WT) and RIG-I(I875A) cells (Fig. 2E). In contrast to a previous study (Ren et al., 2019), we could not observe an influence of N668 on discrimination of p-dsRNA using 24mer dsRNA (Suppl. Fig. 2B). With the ligands used in our study, the RIG-I(N668A) mutant responded to p-dsRNA, OH-dsRNA and ppp-dsRNA in a manner similar to RIG-I(WT).

To confirm that the bulky side chain of I875 prevents CTD-5’p-dsRNA binding, we performed binding studies using surface plasmon resonance (SPR). WT CTD and I875A CTD had *K*_D_ values in the nanomolar range for 5’ppp-dsRNA while no binding was detected for ssRNA (Fig. 2F). WT CTD binding to 5’OH-dsRNA was 15 times stronger than its binding to 5’p-dsRNA. Binding of 5’p-dsRNA to I875A CTD was 10 times stronger compared to WT CTD (Fig. 2E). Unexpectedly, I875A mutation slightly reduced 5’ppp-dsRNA binding and considerably impaired binding of 5’OH to the RIG-I(I875A) CTD without changing the EC50 of OH-dsRNA stimulated full-length RIG-I(I875A) compared to RIG-I(WT). To corroborate the binding results, we determined two crystal structures: A 1.89 Å co-crystal structure of the human RIG-I(WT) CTD with 5’OH-dsRNA (7BAH) and a 3.4 Å structure of the RIG-I(I875A) CTD mutant in complex with 5’p-dsRNA (7BAI; Suppl. Fig. 3A). Of note, we were not able to generate crystals of the RIG-I(WT) CTD with 5’p-dsRNA. Overall, the two 7BAH and 7BAI structures are very similar (Figure 3A). While the resolution of the 5’p-dsRNA structure was low (3.4 Å), a simulated annealing omit map unequivocally confirmed the presence of the 5’p group in the binding pocket of the RIG-I(I875A) CTD (Suppl. Fig. 3B). Fig. 3B shows a comparison of the RIG-I(WT) CTD/5’pp-dsRNA co-crystal structure (3NCU; (Wang et al., 2010)) that of RIG-I(WT) CTD/5’OH-dsRNA (7BAH, this work) and of a previously determined RIG-I(WT) CTD/5’OH-dsRNA structure with a different RNA sequence and length (3OG8 (Lu et al., 2010, 2011)). The 5’pp-dsRNA is held in position by numerous polar and ionic interactions between both the α-(K861, K888) and β-phosphate (H847, K858, K861). Due to the absence of phosphate groups, 5’OH-dsRNA cannot interact with these residues but instead sinks deeper into the binding site. By modeling a 5’p onto the 5’OH-dsRNA structure, we determined that the 5’p would sterically clash with both I875 and G874 in this configuration (Fig. 3C). The lack of interactions with the β-phosphate and the steric repulsion from I875 and G874 thus explain why 5’p does not productively bind to the pocket of the WT CTD. This was corroborated by our RIG-I(I875A) CTD/5’p-dsRNA structure (Figure 3D). Here, the less bulky side chain (of A875) together with numerous polar and ionic interactions with residues K888, G874, K861 and K858 allow the 5’p to be accommodated in the dsRNA binding site without steric clashes. Compared to the 5’pp complex structure, some sidechain rearrangements were observed as shown in Figure 3D. Here, it should be noted that the 5’p has an additional negative charge at the position of the in itself neutral phospho-anhydride bond of the 5’pp, which is reflected in an additional ionic interaction with the amino group of K858 (Fig. 3, Suppl. Fig. 3C). Moreover, due to the low resolution of the 5’p I875A structure, it is not possible to study the influence of the solvent molecules that are involved in the interaction.

**Figure 3:**
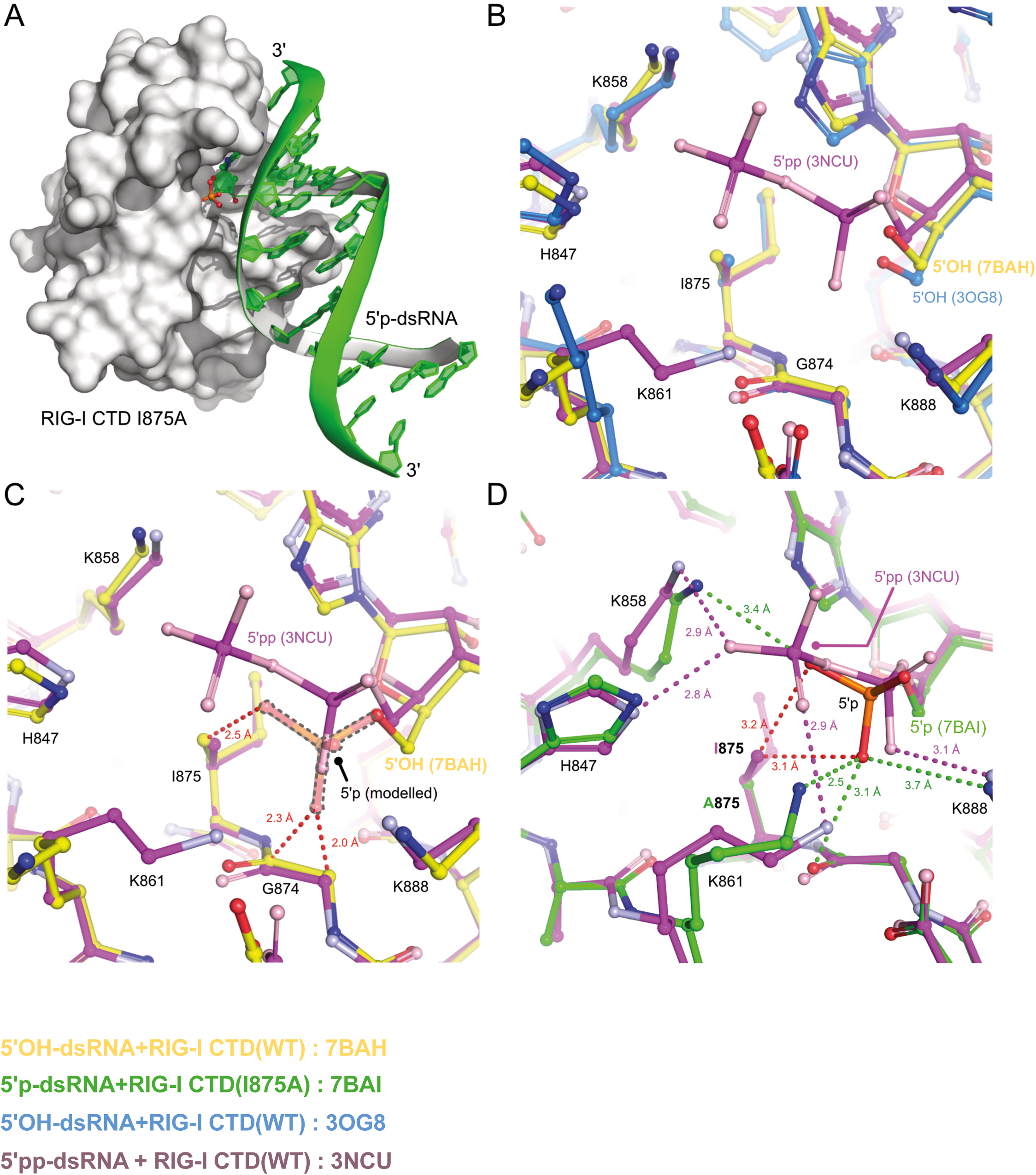
5’p-dsRNA exclusion by the RIG-I CTD. A) Overall structure of the RIG-I-CTD I875A/5’p-dsRNA complex. The CTD is shown as a white surface. The RNA is depicted in green with the terminal monophosphorylated base as a ball-and-stick model. The phosphate atom is colored orange and the phosphate oxygens are colored red. B) Shown is the structural alignment of three ball-and-stick models of RIG-I CTD crystal structures: the 5’-OH-dsRNA/CTD WT complex structure (7BAH, this work, in yellow), the 5’pp-dsRNA/CTD WT structure (3NCU, in magenta), and a previously solved 5’-OH-dsRNA/CTD WT structure (3OG8, in blue) with a different RNA sequence. C) Modelling a monophosphate onto our 5’OH-structure (7BAH, translucent) reveals that I875 and G874 together prevent binding of a monophosphorylated (5’p)-dsRNA. The red dashed lines indicate clashes between the modelled phosphate group and the CTD. D) Crystal structure of the CTD I875A 5’p-dsRNA structure (7BAI, green ball-and-stick model) in comparison to the 5’pp-dsRNA/CTD WT structure (3NCU, magenta ball-and-sticks). Dashed lines indicate distances measured within the two structures. The red dashed lines indicate the very close contacts that would result from a monophosphate bound to the wild-type structure in the same configuration as found in the I875A mutant. RNA sequences: Table S1.

### Tolerance of 5’p prevents recognition of endogenous mtRNA

Since the most abundantly expressed endogenous RNAs (rRNA and tRNA) are processed into molecules with 5’p termini, we speculated that the exclusion of structured 5’p-RNA by WT RIG-I is a mechanism to differentiate between self and non-self RNA. To test the degree of stimulation of RIG-I by endogenous RNA, expression of RIG-I(WT), RIG-I(I875A), and, as a control, RIG-I(H830A), a mutant which senses N1-2’O-methylated 5’pppRNA (Schuberth-Wagner et al., 2015), were induced in RIG-I/MDA5 deficient (RM^-/-^) T-REx-293 cells using an integrated doxycycline inducible locus. Cell-intrinsic IFN signaling was measured via IFIT1 mRNA transcription and IRF3 phosphorylation levels 72h after doxycycline treatment and without the addition of exogenous ligands (Fig. 4A). Of note, after doxycycline treatment, the H830A and I875A mutants were expressed at the same levels as WT RIG-I (Suppl. Fig. 4A, B). Moreover, as expected, long-term expression of RIG-I(H830A) caused an increase in immune activation compared to RIG-I(WT) (Fig. 4A, Suppl. Fig. 4D). However, this effect was even stronger in cells expressing RIG-I(I875A), which was strongly indicative of the presence of endogenous, monophosphorylated, base-paired RNA structures capable of stimulating RIG-I in the absence of a mechanism to actively exclude 5’p from the CTD. Intriguingly, A549 cells expressing the RIG-I(I875A) mutant from the endogenous loci spontaneously secreted CXCL10 and induced an ISG signature including IFNB1, demonstrating the biologic relevance of I875 to prevent RIG-I activation at baseline, homeostatic conditions (Fig. 4B,C).

**Figure 4:**
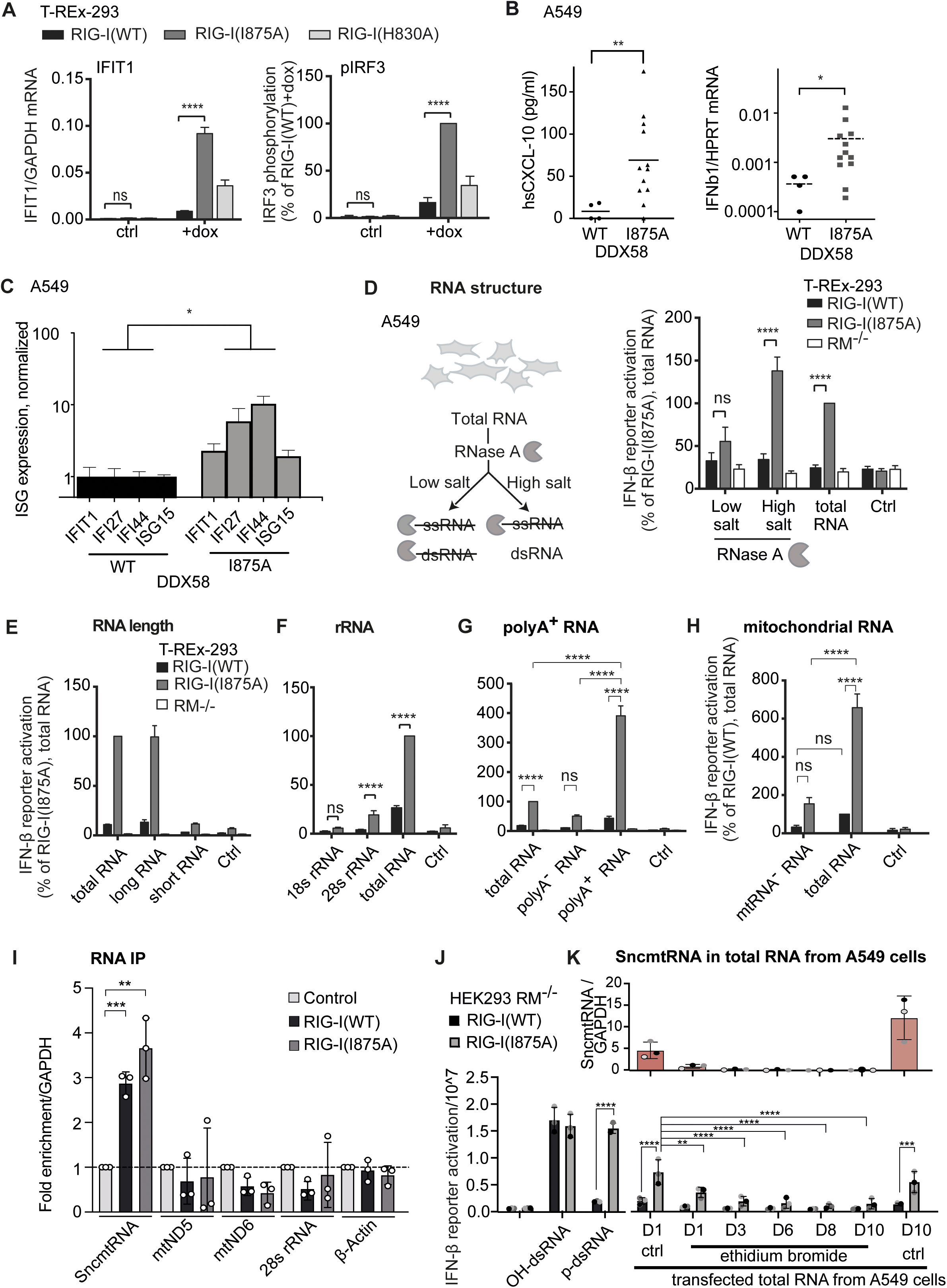
The 5’p tolerance mechanism prevents recognition of endogenous mtRNA by RIG-I. A) RIG-I(WT), RIG-I(I875A) or RIG-I(H830A) expression was induced for 72h with 10 pg/ml doxycycline (+dox) in HEK T-REx RM^-/-^ cells in the absence of exogenous RIG-I stimuli. Left panel: RNA was extracted and mRNA expression of IFIT1 was analyzed by qPCR and normalized to GAPDH (n=3-4, mean + SEM, two-way ANOVA, Tukey’s post-test). Right panel: Phosphorylated IRF3 protein was analyzed by western blot and normalized to β-Actin (n = 4, mean + SEM, two-way ANOVA, Tukey’s post-test). B) Left panel: CXCL-10 levels in WT and DDX58^(I875A/I875/-)^ mutant A549 cells (3 clones, 4 datapoints per clone), 24 h after medium exchange. Shown are cumulated results from two independent experiments. **p<0.01, Welch’s t-test. Right panel: Spontaneous IFN-ß1 mRNA expression in WT and DDX58^(I875A/I875/-)^ mutant A549 cells (3 clones) measured by qPCR. RNA was isolated from growing cultures twice per week. Shown are cumulated data from 4 timepoints, individual values and mean (dotted line). *p<0.05, Welch’s t-test. C) Spontaneous ISG signature of WT and DDX58^(I875A/I875/-)^ mutant A549 cells (3 clones) measured by qPCR. RNA was isolated from growing cultures twice per week. Shown are means + SEM of cumulated data from 4 timepoints per culture, normalized for the average of expression in WT. *p<0.05, Mann-Whitney test. D) Total RNA from A549 cells was treated with RNase in low-salt (50 mM NaCl) or high-salt (300 mM NaCl) conditions or left untreated. RNA (1 µg/ml) was transfected into HEK T-REx RM^-/-^ cells expressing RIG-I(WT), RIG-I(I875A) or no RIG-I (RM^-/-^). IFN-β reporter activation was determined 20 h after RNA transfection and is depicted as percentage of the signal of RIG-I(I875A) stimulated by non-treated total RNA (n = 5, mean + SEM, two-way ANOVA, Tukey’s post test). E) Total RNA was separated into a short RNA(18-200nt) and a long RNA(>200nt) fraction. HEK T-REex RM^-/-^ cells expressing either RIG-I(WT), RIG-I(I875A) or no RIG-I (RM^-/-^) were stimulated with the indicated RNA (1 µg/ml). IFN-β reporter activation was determined 20h after RNA transfection and is depicted as percentage of the signal of RIG-I(I875A), stimulated with total RNA (n = 4, mean + SD). F) Total RNA was separated on an agarose gel and 18s rRNA and 28s rRNA were extracted from the gel. HEK T-REx RM^-/-^ cells expressing RIG-I(WT) or RIG-I(I875A) were transfected with total RNA and the extracted 18S rRNA and 28S rRNA (1 µg/ml). IFN-β reporter activation was determined 20h after RNA transfection and is depicted as percentage of the signal of RIG-I(I875A), stimulated with total RNA (n = 4, mean + SD, two-way ANOVA, Sidak’s post-test). G) Total RNA was fractionated into a polyA^+^ and polyA^-^ fraction using polyA Beads. Fractionated RNA and total RNA (1 µg/ml) was transfected into HEK T-REx RM^-/-^ cells expressing RIG-I(WT), RIG-I(I875A) or no RIG-I (RM^-/-^). IFN-β reporter activation was determined 20h after RNA transfection and is depicted as percentage of the signal of RIG-I(I875A) stimulated by total RNA, (n = 5, mean + SEM, two-way ANOVA, Tukey’s post test). H) Total RNA was extracted using TRIzol from A549 cells that were cultivated in ctrl medium (total RNA) or ethidium bromide containing medium (mtRNA^-^) for at least 7 days. HEK T-REx RM^-/-^ cells expressing RIG-I(WT) or RIG-I(I875A) were transfected with the RNA (1 µg/ml). IFN-β reporter activation was determined 20 h after RNA transfection and is depicted as percentage of the signal of RIG-I(WT) stimulated by total RNA (n = 3 (RIG-I(WT) and RIG-I(I875A)), mean + SEM, two-way ANOVA, Tukey’s post test). I) FLAG-RIG-I(WT) or FLAG-RIG-I(I875A) were overexpressed in HEK293FT cells. 24 h after transfection cells were lyzed, FLAG-RIG-I proteins immunoprecipitated using anti-FLAG magnetic beads and bound RNA was extracted and used for qPCR (n = 3 (RIG-I(WT) and RIG-I(I875A)), mean + SEM, two-way ANOVA, Tukey’s post test). J) 24mer OH-dsRNA, 24mer p-dsRNA or cellular RNA isolated from 1,3,6,8 or 10 days EtBr- or un-treated (ctrl) A549 cells was transfected into RIG-I(WT) or RIG-I(I875A) expressing HEK293 cells and RIG-I stimulation monitored by IFN-β reporter activation. K) qPCR of SncmtRNA in total RNA from A549 cells used for stimulation in (J). Statistical significance is indicated as follows: ** (p < 0.01), *** (p < 0.001), **** (p < 0.0001), ns: not significant. Dox: Doxycycline.

To identify the endogenous RNA species causing the observed RIG-I(I875A) activation, whole-cell RNA was isolated and transfected into the HEK T-REx cell lines. As expected, total RNA induced stronger IFNB promoter reporter activation in RIG-I(I875A) cells compared to RIG-I(WT) (Suppl. Fig. 4E). To better characterize these endogenous RNA ligands, selective RNase A digestion was then employed: RNase A digests both ssRNA and dsRNA at low-salt concentrations (up to 100 mM NaCl), while at high-salt concentrations (>300 mM NaCl), it selectively digests ssRNA but not dsRNA. Whole-cell RNA lysate digested with RNase A in a high-salt buffer triggered the same level of IFNB promoter reporter activation as untreated RNA in HEK T-REx cells. However, total cellular RNA treated with RNase A under low-salt conditions reduced the immune stimulatory potential of the RNA significantly compared to untreated RNA, indicating that the stimulatory RNA consisted of base-paired structures (Fig. 4D). Moreover, we detected RIG-I(I875A)-stimulating RNA within the fraction of long (>200nt) but not short RNAs (18-200 nt) (Fig. 4E). rRNAs and tRNAs both form complex secondary structures with significant regions of base-pairing. Thus, to identify the specific stimulatory species within the total cellular RNA pool, RNA was fractionated and purified. tRNA isolated from yeast did not activate an immune response in either RIG-I(WT) or RIG-I(I875A) cells (Suppl. Fig. 4F). To specifically test the stimulatory potential of 18S and 28S rRNA, total RNA was separated on an agarose gel and the 18S and 28S bands were excised and extracted. While stimulation with 18S rRNA had no significant effect on RIG-I(WT) or RIG-I(I875A) cells, 28S rRNA stimulation led to significant IFN-β induction in RIG-I(I875A) cells (Fig. 4F). Nonetheless, the observed IFN-β induction was five times lower than stimulation with total RNA (Fig. 4F). To exclude that this discrepancy resulted from the fractionation method used, a second rRNA extraction method was applied. Specific locked-nucleic acid (LNA) probes against 18S and 28S rRNA were used to pull-down rRNA from the total cellular RNA pool. Both rRNA isolated by this method (rRNA^+^) and the resultant fraction without rRNA (rRNA^-^) activated RIG-I(I875A) cells at a similar level, which was 6-fold higher than in RIG-I(WT) cells (Suppl. Fig. 4G). While these data demonstrate that rRNA can contribute to the immune activation potential of total cellular RNA extracts, it also indicates that they are not the only immune-activating species within this RNA. Thus, we then examined the 3’-polyadenylated fraction of total RNA using polydT beads for RNA purification. mRNAs are the main polyadenylated RNA species, but rRNAs containing polyA tails have also been described (Slomovic et al., 2006). Like tRNAs and rRNAs, mRNAs are single-stranded but can form secondary structures with stretches of base-pairing. Of note, mRNAs have a cap1 structure (not a 5’p) and thus would not be expected to stimulate RIG-I(WT) or I875A. Surprisingly, the fraction containing the polyA RNA robustly induced IFN-β reporter activation in RIG-I(I875A) cells, as compared to total RNA (Fig. 4G). Accordingly, the fraction that had been depleted of polyA RNA demonstrated less IFN-β reporter activation in comparison to total RNA (Fig. 4G). Therefore, polyA RNA seems to be the main activating species of RIG-I(I875A).

Although mRNA has a cap1 structure, thereby preventing activation of wild-type RIG-I and RIG-I(I875A), mitochondrial mRNAs (mt-mRNAs) are both uncapped and are reported to possess a stable 3’-polyA tail (Fernandez-Silva et al., 2003; Ojala et al., 1981). Indeed, we could verify this observation via qPCR on the polyA RNA fraction, which revealed the presence of the mt-mRNAs ND1 and ND5 (Suppl. Fig. 4H). Human mitochondria possess multiple copies of their 16.6-kb circular dsDNA genome that is bidirectionally transcribed and encodes 11 mt-mRNAs, two mt-rRNAs and 22 mt-tRNAs (Anderson et al., 1981; Mai et al., 2017). The mt-tRNAs are excised from a long polycistronic precursor RNA by RNase P and Z, leading to the release of the mt-mRNAs, which are located between the tRNAs on the mitochondrial genomic DNA strands (Rossmanith, 2012). RNase Z cleavage results in 3’OH and 5’P termini, thus generating 5’P mt-mRNAs, which are polyadenylated by a human nuclear-encoded mitochondrial poly(A) polymerase (Mai et al., 2017). Villegas et al. (Villegas et al., 2007) identified mitochondria derived polyadenylated long non-coding RNAs (e.g. *SncmtRNA;* DQ386868.1) forming a hairpin with a long (200-800 bp) stem structure which was localized in the cytoplasm and nucleus of proliferating cells. Furthermore, the occurrence of double-stranded mtRNA that activates MDA5 and PKR has already been demonstrated (Dhir et al., 2018; Kim et al., 2018).

To determine the contribution of mtRNA to the activation of RIG-I(I875A), we used ethidium bromide (EtBr) to deplete mtRNA levels. Long-term incubation of cells with low doses of EtBr causes a loss of mt-DNA (and thus mtRNA) due to a decrease in its replication rate (King and Attardi, 1996). Since HEK T-REx cells did not survive EtBr treatment (data not shown), A549 cells were used instead for mtRNA depletion. After incubation for at least seven days in medium containing EtBr, whole-cell RNA was extracted from these cells (mtRNA^-^) and from cells that were cultivated in a control medium (mtRNA^+^). The expression of rRNAs and nuclear-encoded mRNAs (GAPDH) did not change between RNA from EtBr-treated and mtRNA^+^ A549 cells (Suppl. Fig. 4I). However, expression of mt-mRNAs (ND1, ND5 and SncmtRNA) was greatly reduced in cells treated with EtBr, confirming the EtBr-mediated depletion of mt-DNA/RNA (Suppl. Fig. 4I, J). RIG-I(WT) and RIG-I(I875A) cells were then transfected with the RNA from non-treated (mtRNA^+^) and EtBr-treated cells (mtRNA^-^). As expected, RNA from the mtRNA^+^ cells led to an elevated immune response in RIG-I(I875A) expressing cells. However, this effect was completely abrogated by previous EtBr-treatment (Fig. 4G). To identify mtRNA interacting with RIG-I, we expressed FLAG-RIG-I(WT) or FLAG-RIG-I(I875A) in HEK293T cells, immunoprecipitated FLAG-tagged proteins including bound RNAs from transfected cells or mock and analyzed candidate RNAs (28s rRNA, Actin, ND5, ND6, SncmtRNA and GAPDH) by qPCR. Intriguingly, among candidate RNAs only SncmtRNA was found to be enriched in pull-downs of FLAG-RIG-I(WT) or FLAG-RIG-I(I875A) expressing cells in comparison to mock (Fig. 4I), indicating that RIG-I binds SncmtRNA but not the mtRNAs ND5, ND6 or ribosomal 28s rRNA. To further correlate mtRNA with RIG-I(I875A) stimulating activity in total RNA, we performed a time course of EtBr treatment of A549 cells and isolated total RNA from 1,3,6,8 or 10 days EtBr-treated or -untreated cells. The isolated total RNA was transfected into RIG-I(WT) or RIG-I(I875A) expressing HEK293 cells, and RIG-I stimulation was monitored by IFN-β reporter activation (Fig. 4J). RIG-I activation correlated well with SncmtRNA levels over different incubation times (Fig. 4K), further supporting the notion that mtRNA is crucial for the observed RIG-I activation. To further validate that monophosphorylated SncmtRNA is a potential RIG-I stimulating ligand, we synthesized sense and antisense strands by *in vitro* transcription which, after hybridization, simulate the structure of SncmtRNA. Since an IVT reaction cannot start with a uridine, we designed two different sense strands (1 or 2) that either start 10 nt behind the original start or have 3 additional nt upstream (5’) of the uridine complementary to the antisense strand, yielding a 5’base-paired structure with 3’overhang, as found in the original sequence (Fig. 5A). 5’ppp was digested with polyphosphatase leaving a 5’monophorylated end. Here, only RIG-I(I875A) expressing cells showed a substantial immune response to monophorsporylated IVT SncmtRNA (Fig. 5B).

**Figure 5:**
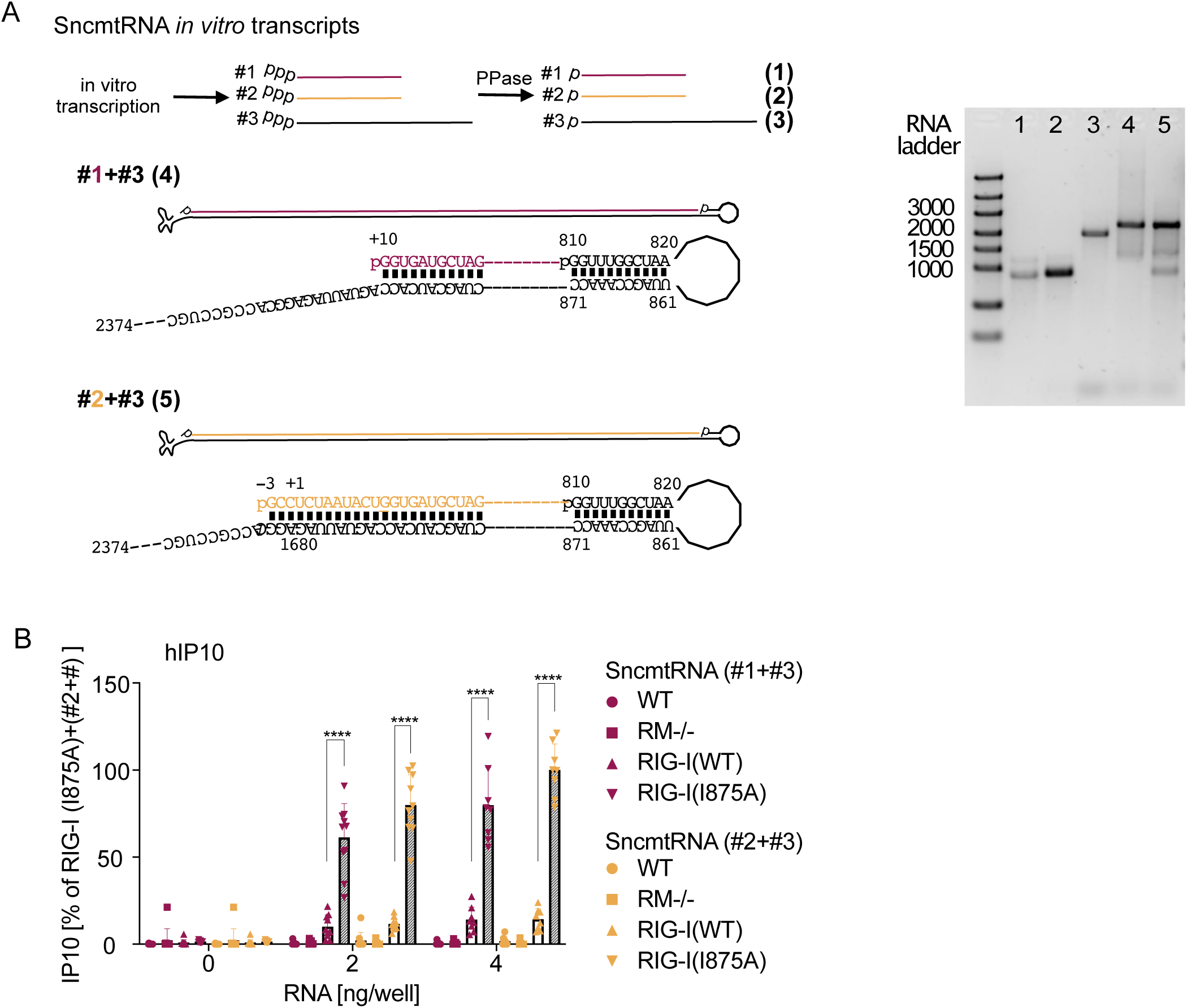
RIG-I(I875A) recognizes monophosphorylated SncmtRNA *in vitro* transcripts. A) Scheme of constructs mimicking the human SncmtRNA (DQ386868.1). Indicated parts of the SncmtRNA gene (DQ386868.1: #1: +10 - 809; #2: −3 - 809: #3: 810-2374) were amplified, fused to a 5’T7-promoter and generated by *in vitro* transcription. 5’triphosphate(ppp) ends were digested by 5’polyphosphatase (PPase) which left a 5’monophosphate (p). #1+#3 or #2+#3 were annealed and B) transfected into WT, RIG-I/MDA5 deficient (RM^-/-^) or RM^-/-^ Flp-In™ T- REx™ 293 cells expressing RIG-I(WT) or RIG-I(I875A) upon addition of 0.1 ug/ml doxicycline. IP10 in superantants was determined by ELISA 20 h after stimulation (n=10). C) Agarose gel of indicated RNAs: #1, #2, #3 and hybridized #1+#3 and #2+#3. IVT template cloning primer sequences: Table S4

Altogether, these results demonstrate that, within total RNA, polyadenylated mtRNA is the main species stimulating RIG-I(I875A) and that 28s rRNA has only a minimal to negligible role in the endogenous stimulation of RIG-I. Thus, we conclude that if RIG-I WT was not able to exclude 5’p-RNA from its binding site, mtRNA and particularly SncmtRNA would induce constitutive activation of RIG-I as observed for RIG-I(I875A) expressing cells, leading to chronic autoinflammation.

## DISCUSSION

Most cytosolic antiviral innate immune receptors (PKR, OAS, RLR) recognize dsRNA (reviewed in (Schlee and Hartmann, 2016)). Originally, it was presumed that dsRNA is a marker for non-self, as it was thought to only occur during viral replication and be otherwise absent from vertebrate cells (Weber et al., 2006). However, some recent studies, including our own, have exhibited the necessity for additional levels of control to prevent recognition of self-dsRNA by these receptors: ADAR1 modifies dsRNA stretches within polymerase II transcripts to abrogate base pairing that would otherwise activate PKR and MDA5 signaling (Pestal et al., 2015; Pfaller et al., 2018; Tang et al., 2021); the RNA-degrading enzyme, SKIV2L, part of the RNA exosome, limits activation of RLR by endogenous RNA (Eckard et al., 2014); the RNA helicase SUV3 and the RNA degrading enzyme polynucleotide phosphorylase (PNPase) restrict the levels of mitochondrial dsRNA that can stimulate MDA5 (Dhir et al., 2018). Constitutive activation of RIG-I or MDA5 by endogenous RNA would likely have a phenotype similar to RIG-I or MDA5 gain-of-function mutations, which have been shown to cause autoimmune diseases like Singleton-Merten syndrome or Aicardi-Goutières syndrome (Lassig et al., 2018; Rehwinkel and Gack, 2020). While MDA5 recognize very long dsRNA (>1000bp) (Kato et al., 2008), RIG-I can detect short stretches of base-paired RNA (>20bp) (Marques et al., 2006; Schlee et al., 2009), the incidental occurrence of which is virtually unavoidable. Therefore, strict tolerance mechanisms for self-RNA are especially important for RIG-I activation. Our group previously discovered that a single amino acid, H830, in the RIG-I CTD is crucial to avoid activation by cap N1-2’O-methylated RNA (Schuberth-Wagner et al., 2015). This mechanism mediates tolerance to endogenous cap1-RNA. In this study, we found that the presence of a 5’-monophosphate terminus is sufficient to nearly completely prevent activation of RIG-I, even by long dsRNA. This finding clearly demonstrates that an entirely 5’terminus-independent RIG-I activation by long (>200bp) dsRNA as suggested in a study comparing 5’ppp-dsRNA and 5’OH-dsRNA (Binder et al., 2011) does not exist. By systematically testing different endogenous RNA species, we could show that 5’p-discrimination has a strong impact on the recognition of mitochondrial RNA. Although tRNA and rRNA represent the largest fraction of 5’p-bearing RNA in the cytosol (up to 95%), our data, using depletion and enrichment of rRNA and monitoring of 28s rRNA specifically bound to FLAG-RIG-I pull-downs, indicate a rather subordinate role for (28s) rRNA in RIG-I(I875A) activation and no immune activation at all by tRNA. Here, it is plausible that the extensive internal post-transcriptional modifications of rRNAs and tRNAs prevent RIG-I binding and activation (Hornung et al., 2006). Moreover, endogenous vault RNA, a short, structured 5’p-RNA, was described to stimulate RIG-I during viral infections if enriched as non-processed 5’ppp-RNA (Zhao et al., 2018). However, we did not observe RIG-I(I875A) activation for the short RNA fraction (18-200 nt) from total RNA. In contrast, depletion of either polyadenylated RNA or mitochondrial RNA from isolated total RNA resulted in an almost complete loss of the RIG-I(I875A) signal. Of note, all mitochondrial RNA species with the exception of mt-tRNA are 3’polyadenylated (Slomovic et al., 2005). While nuclear-transcribed mRNA is capped at the 5’-end and therefore excluded from RIG-I binding, mt-mRNA lacks a 5’-cap structure. mtRNA is polycistronic and cleaved by RNase P and an RNase Z after transcription to form 5’p-mt-tRNA and 5’p-mt-mRNA. Because transcription of the circular mitochondrial genome is bidirectional, mtRNAs from opposite directions of transcription can form 5’p-dsRNA (Kim et al., 2018). The 5’p termini protect these mt-dsRNA species from wildtype RIG-I but not RIG-I(I875A) recognition. Additionally, long non-coding (lnc)-mtRNA, such as “SncmtRNA” or “ASncmtRNA”, are generated by post-transcriptional events leading to the fusion of 16s mt-rRNA with complementary mtRNA and harbor a long stem (>800bp) with 3’overhang. Such structures would be amenable targets for RIG-I if 5’p did not prevent RIG-I activation (Schlee et al., 2009) but not for MDA5, which requires more than 1000bp (Kato et al., 2008). SncmtRNA is also polyadenylated and was found in the nucleus and cytosol of proliferating cells (Landerer et al., 2011; Villegas et al., 2007), indicating an active cytosolic transport and function in cell maintenance. Strikingly, we observed specific interaction of RIG-I and RIG-I(I875A) with SncmtRNA but not with 28s rRNA, housekeeping genes (GAPDH, Actin) or the mtRNAs coding for ND5 or ND6 .

In line with our findings, a previous study has already demonstrated that most cellular dsRNA originates from mtRNA (Dhir et al., 2018). Of note, Kim et al. demonstrated that the cytosolic dsRNA binding receptor PKR is constantly activated by endogenous mtRNA (Kim et al., 2018), while Dhir et al. showed that MDA5 detects mtRNA if the enzymes involved in mtRNA degradation have been depleted (Dhir et al., 2018). Since depletion of mtRNA from total RNA completely removed its RIG-I(I875A) stimulatory activity, we conclude that mt-dsRNA is the main RNA species activating RIG-I(I875A). While it is unclear if mt-mRNA is released at sufficient levels to activate RIG-I during cell maintenance, lnc-mtRNAs, which are known to reside in the cytosol and nucleus, are the more plausible RIG-I targets. Our data suggest that RIG-I would be constantly activated by mtRNA --at least in proliferating cells--if it lacked the ability to discriminate between 5’p and 5’OH-dsRNA. Interestingly, viral RNA can also bear a 5’p “tolerance motif”. The (+)ssRNA genome of HCV is cleaved by RNase P, the same nuclease that also cleaves tRNA precursors and produces 5’p ends (Nadal et al., 2002). Some (-)ssRNA viruses, including Crimean-Congo hemorrhagic fever virus (CCHTV), Hantaan virus (HTNV), and Borna disease virus (BDV), cleave their 5’ppp-end to leave a 5’p-end (Habjan et al., 2008). Thus, it is not only the absence of 5’ppp but also the presence of a 5’p contributes to the repression of the RIG-I mediated detection of these viruses, which is in line with the exploitation of other self tolerance motifs by other RNA viruses to avoid innate immune activation, e.g. N1-2’O-methylation of yellow fever RNA or cap-snatching by influenza.

Binding and activation assays revealed that 5’p-dsRNA exclusion results from the the bulky methyl groups of I875 within the triphosphate binding pocket of RIG-I. Consequently, mutation of I875 to alanine facilitated binding and enabled activation of RIG-I by 5’p-dsRNA. RNAs with more than one phosphate, namely 5’pp-dsRNA and 5’ppp-dsRNA, are still able to bind RIG-I in the presence of I875. However, in this case, the additional, strong interactions between the amino groups of K861, H849, and K858 with the β-phosphate stabilize the ligand in a conformation that does not clash with I875. If 5’p-dsRNA were accommodated into the CTD in the same binding mode as 5’OH-dsRNA, as observed in 7BAI, the 5’p would clash with both I875 and the peptide backbone at G874. The proximity of I875 to the α-phosphate or the clash of a modeled 5’p with G874 can also be observed in structures of human RIG-I(WT) CTD with 5’ppp-dsRNA (3lrn) or 5’OH-dsRNA (3og8) with a different core sequence and length (16mer) (Lu et al., 2010, 2011). Thus far, we have no explanation why I875A mutation impairs binding of 5’OH-dsRNA to the RIG-I(I875A) CTD and why this does not have consequences for full-length RIG-I(I875A) activation by the same ligand.

The group of Fujita (Takahasi et al., 2008) described RIG-I stimulation by monophosphorylated dsRNA (5’p at one end, 5’OH at the other) in L929 cells. However, our data demonstrated that the presence of a 5’p at both ends of the same dsRNA sequence abolished the activation of RIG-I, thus strongly indicating that it was the 5’OH-end rather than the 5’p-end of that sequence that induced RIG-I activation. Additionally, the RNA-stabilizing effect that led to RIG-I stimulation in L929 cells, a murine fibroblast cell line, might not be present or play a role in the cells that we used in our study.

Ren et al. also described inhibition of RIG-I binding to 5’p-dsRNA when compared to 5’OH-dsRNA (Ren et al., 2019) but claimed a critical “gatekeeper” role for N668 in the RIG-I helicase domain. However, considering that the first high affinity contact with the incoming RNA is made with the free CTD (before it has formed a complex with the helicase domain), it is unclear how amino acids outside of the CTD could plausibly take on the role of a “gatekeeper”. Additionally, Ren et al. found that the binding affinities of *all* tested RNAs (ppp-RNA, pp-RNA, p-RNA and OH-RNA) were decreased for RIG-I(N668A) compared to RIG-I(WT), and the N668A mutation in the helicase domain did not show a specific activity effect: It led not only to increased interferon induction by 5’p-dsRNA but also by 5’OH-dsRNA (Ren et al., 2019). Using 24mer dsRNA we could not detect a considerable difference between RIG-I(WT) and RIG-I(N668A) concerning recognition of p-dsRNA, OH-dsRNA or ppp-hpRNA. We can only speculate that the observed effect of the Ren group is due to the use of very short hairpin RNA and that the fourfold enhancement of *both* p-dsRNA and OH-dsRNA recognition observed for RIG-I(N668A) is due to a changed helicase unwinding or translocation activity of RIG-I that influences RIG-I activation or the stability of the RNA/RIG-I complex since N668 is close to T667, a residue critically involved in RIG-I translocation on the RNA (Devarkar et al., 2018). Of note, in contrast to I875, N668 is not a highly conserved amino acid in RIG-I (in vertebrates N can be replaced by D, S or T). The lack of RIG-I(WT)-mediated type-I-IFN induction by 5’OH-dsRNA in the Ren paper is most probably due to the usage of very short ligands (12bp) (Ren et al., 2019). Ligands at this length cannot activate RIG-I at moderate ppp-dsRNA concentrations (Schlee et al., 2009), rendering it even more difficult to detect the signal induced by weaker ligands in a biological context. In contrast, we showed that I875A mutation in the CTD selectively allows 5’p-dsRNA binding and RIG-I activation, demonstrating the role of I875 in preventing 5’p-dsRNA binding. Furthermore, activation of other RNAs (5’OH-dsRNA, 5’ppp-dsRNA, or RNA with overhangs) was unchanged between RIG-I(WT) and the RIG-I(I875A) mutant, highlighting the specificity for the steric exclusion of 5’p-dsRNA.

Altogether, our study identifies a novel tolerance mechanism within the binding pocket of RIG-I that helps ensure that endogenous mitochondrial RNA does not lead to self-activation of the innate immune system. As, to our knowledge, RIG-I is expressed in almost all nucleated cells, inappropriate activation by mitochondrial RNA would have potentially devastating effects on the host. In line with previous studies (Dhir et al., 2018; Kim et al., 2018), our study identifies endosymbiotic mitochondria as a source of unwanted, potentially stimulatory nucleic acids, underscoring the effort that the host cell has had to undertake to accommodate its ancient guest.

## ACKNOWLEDGEMENTS

We thank Meghan Campbell for her critical reading of the manuscript.

This study was funded by the Deutsche Forschungsgemeinschaft (DFG, German Research Foundation) under Germany’s Excellence Strategy – EXC2151 – 390873048 of which EB, KP, GH, MG and MS are members. It was also supported by other grants of the DFG, including TRR237 (EB, TZ, KP, GH, MS), SFB670 (EB, GH, MS), SFB704 (GH), GRK2168 (EB, MS) and DFG SCHL1930/1-2. EB and MS received financial support from BONFOR (University of Bonn). This work is part of the PhD thesis of AKdR at the University of Bonn and of the Bachelor thesis of Bastian Putschli at the Johannes Gutenberg University Mainz.

## AUTHOR CONTRIBUTIONS STATEMENT

AkdR, KA, DF, TZ, BP, KG, AG, SW, DH, KC, AK, CH CSW performed the experiments. AkdR, KA, KG, DH, KC, AK, SW, EB, KP, JL, MG and MS conceived experiments. AkdR, KG, EB, KP, GH, GrH, MG and MS analyzed and interpreted data and wrote the manuscript. All authors revised the manuscript.

## MATERIALS AND METHODS

### LEAD CONTACT AND MATERIALS AVAILABILITY

Further information and requests for resources and reagents should be directed to and will be fulfilled by the Lead Contact, Martin Schlee. Materials will be made available via material transfer agreement (MTA).

### DATA AND CODE AVAILABILITY

The crystal structures generated during this study are available at Protein Data Bank (7BAH, 7BAI).

### EXPERIMENTAL MODEL AND SUBJECT DETAILS

#### Primary cells

Human PBMCs were isolated from buffy coats of healthy human donors and cultivated in RPMI 1640 medium (10% FCS, 100 U/ml penicillin, 100 mg/ml streptomycin) at 37 °C.

#### Cell lines

A549 cells were cultivated in DMEM (10% FCS, 100 U/ml penicillin, 100 mg/ml streptomycin) at 37 °C. HEK Flp-In^™^ T-REx^™^ 293 were cultivated in DMEM (10% FCS, 100 U/ml penicillin, 100 mg/ml streptomycin) at 37 °C.

### METHODS DETAILS

#### Cell culture

Human PBMCs were isolated from whole human blood of voluntary, healthy donors by Ficoll-Hypaque density gradient centrifugation (Merck, Darmstadt, Germany). For stimulation experiments, 4 x 10^5^ PBMCs were cultured in 96-well plates. The PBMC studies were approved by the local ethics committee (Ethikkommission der Medizinischen Fakultät Bonn) according to the ICH-GCP guidelines. To inhibit TLR7/8 activity, cells were pre-incubated with 5 µg/ml chloroquine for 30 min. Cells were kept in RPMI 1640 medium (10 % fetal calf serum (FCS), 100 U/ml penicillin, 100 mg/ml streptomycin).

All transfections were performed using Lipofectamine 2000 (LF2000, Thermo Fisher Scientific, Darmstadt, Germany). The transfection reagent was mixed with OptiMEM and was incubated for 5 min at room temperature (RT). RNA or DNA was mixed with an equal amount of OptiMEM and combined with the OptiMEM-transfection reagent mix. After 20 min of incubation at RT, the transfection mixture was added to the cells.

HEK Flp-In^™^ T-REx^™^ 293 cells (Thermo Fisher Scientific) were kept in DMEM (10% FCS, 100 U/ml penicillin, 100 mg/ml streptomycin). For stimulation experiments, 3.5 x 10^6^ cells were cultured in 96-well plates. Protein and RNA samples were harvested from 24-well plates that had been seeded with 2 x 10^5^ cells. Doxycycline (100 pg/ml or as indicated) was added to the cells in the plate 20 min before the experiments to induce the expression of RIG-I.

#### Generation of stable HEK Flp-In™ T-REx™ 293 expression cell lines

The HEK Flp-In^™^ T-REx^™^ 293 host cells were seeded in 6-well culture plates (1 x 10^6^ cells in 3 ml/well) and allowed to adhere overnight. The following day 700 ng pOG44 plasmid, expressing the Flp recombinase, was transfected simultaneously with 70 ng of pcDNA5/FRT plasmid containing the gene of interest using LF2000. After 48h of incubation, the cells were trypsinized, resuspended in selection media containing 150 µg/ml hygromycin and 10 µg/ml blasticidin and seeded in new 6-well plates. After the selection process, expression of the protein of interest was confirmed by functional assays and western blot.

#### Generation of gene-edited cell lines

HEK Flp-In^™^ T-REx^™^ 293 cells were transfected with LF2000 with 200 ng of a CAS9-gRNA expression plasmid targeting RIG-I (5**-**GGGTCTTCCGGATATAATCC(TGG)-3) or MDA5 (5-CTTCTAGTTAGAGACGTCT TGG (TGG)-3). One day after transfection cells were cultivated in limited dilution to obtain single-cell clones. After expansion knockout (KO) was confirmed by immunoblot, Sanger sequencing and functional testing. To generate A549-RIG-I(I875A) point mutant cell lines, A549 cells were co-electroporated with Cas9-2A-EGFP-gRNA expression plasmid targeting 5’-CAAATGTCTTGTACTTCACA(TGG)-3’ (100 µg/ml) and ssDNA repair oligo 5’-AAGATATTCTGTGCCCGACAGAACTGCAGCCATGACTGGG<u>GA**GCT**C</u>ATGTGAAGTACAAGACATTTGAGATTCCAGTTATAAAAATTGAA-3’ (5µM, I875A mutation bold, generates *SacI* site, underlined), using the Neon Transfection System (Thermo Fisher Scientific). Settings were as recommended by the manufacturer (1230 V, 30 ms, 2 pulses). 18h after electroporation, EGFP-positive cells were enriched by FACS-sort (BD FacsAria III Sorter, Flow Cytometry Core Facility Bonn). Single-cell clones were obtained by limiting dilution and gene-editing was confirmed by *SacI*-digestion and Sanger sequencing of the amplified genomic site (fw 5’-TGGTTCTAACGTTCTCTTTTGTGTG-3’, rev 5’-AGCGATCCATGATTATACCCACT-3’) and functional testing. Clone genotypes are listed in Fig. 2D.

#### Generation of RIG-I mutants

Point-mutations in full-length RIG-I were introduced using PCR amplification of plasmids containing the RIG-I WT gene with mutated primers (see table S3) and gibson assembly. The mutant constructs were confirmed by sequencing. Protein expression of RIG-I mutants in HEK Flp-In^™^ T-REx^™^ 293 cells was compared by immunoblot with wild-type RIG-I using an antibody against RIG-I (Cell signaling, Frankfurt am Main, Germany).

#### Generation of mtDNA - and mtRNA - depleted cells

A549 cells were cultivated in DMEM containing ethidium bromide (50 ng/ml), sodium pyruvate (1 mM), uridine (50 µg/ml) and non-essential amino acid solution (NEAA; 1x) for at least 7 days. Successful mtRNA depletion was measured via gene expression analysis of mt-mRNA (ND1, ND5) using qPCR. Control cells were incubated in medium containing sodium pyruvate (1 mM), uridine (50 µg/ml) and NEAA (1x), but without EtBr.

#### Detection of Cytokines

IFN-α levels were analyzed with commercial ELISA assay kits (Thermo Fisher Scientific) according to the manufacturer’s protocol. The concentration of IFN-α present in the sample was determined by a standard curve, which was determined using known amounts of recombinant cytokines.

#### Dual luciferase assay

HEK Flp-In^™^ T-REx^™^ 293 were seeded in 96-well plates and were allowed to adhere overnight. The cells were then co-transfected with 10 ng Gaussia luciferase reporter plasmid (pGL3-IFNB1-GLuc) and 5 ng control Firefly Luciferase expression plasmid for normalization (pLenti-Ef1α-FLuc) (Gao et al., 2013). After at least 6h, the cells were stimulated with the RNA ligands indicated in the figures. The luciferase assay was performed 20h after stimulation. To measure *Gaussia* luciferase production, as a surrogate for IFN-β promoter activation, 25 µl coelenterazine-solution (1 µg/ml in H_2_O) was added to 25 µl supernatant in a white microplate. Chemiluminescence was immediately measured with the EnVision 2104 Multilabel plate reader (Perkin Elmer, Waltham, USA). To measure Firefly luciferase as a control, the supernatant was aspirated and 40 µl 1 x SAP-solution (0.02 % Saponin, 50 mM tris HCL pH 7.8, 15 mM MgSO_4_, 4 mM EGTA, 10 % Glycerol) was added to the cells and then they were lysed for 20 min at RT on a shaker. Afterwards, 25 µl of lysed cells in SAP were added to 25 µl Firefly luciferase substrate (Luciferin) and luminescence was measured with the EnVision 2104 Multilabel plate reader.

#### Preparation of RNA

Triphosphorylated RNA oligonucleotides were chemically synthesized as described (Goldeck et al., 2014).

5’OH-RNA and 5’p-RNA were ordered from Biomers (RNA sequences are listed in TableS1). 60mer, 100mer, and 200mer RNA were transcribed using *in vitro* transcription (IVT, TranscriptAid™ T7 High Yield Transcription Kit, Thermo Fisher Scientific).

Whole-cell RNA (total RNA) was extracted using TRIzol™ reagent from A549 cells (Thermo Fisher Scientific) according to the manufactureŕs instruction. polyA-RNA extraction: Purification of polyA-containing RNA from the whole-cell RNA was performed with the NucleoTrap® mRNA Kit (Macherey Nagel, Düren, Germany) according to the manufactureŕs instructions.

#### rRNA extraction

rRNA was extracted from whole cell RNA extracts (total RNA) using the RiboMinus™ Kit or by excision and extraction of the 18s and 28s rRNA bands from an agarose gel.

Extraction with the Ribominus™ Human/Mouse Transcriptome Isolation Kit (Thermo Fisher Scientific) was performed according to the manufactureŕs instructions. The rRNA^-^ fraction was isolated using columns while the rRNA^+^ fraction was isolated from the beads using TRIzol extraction.

To extract rRNA using gel electrophoresis, whole cell RNA extracts were run on a 1% agarose gel in RNase free conditions. RNA bands according to the size of the 18s and 28s rRNA were cut from the agarose gel and extraction was performed using the NucleoSpin® Gel and PCR Clean-Up Kit (Macherey-Nagel). The extraction was performed according to the manufactureŕs protocol for RNA extractions from agarose gels using the buffer NTC (Macherey Nagel).

#### Small RNA extraction

Small RNA (<200 nucleotides) was extracted from whole cell RNA extracts (total RNA) using the miRNeasy Mini Kit and the RNeasy MiniElute Cleanup Kit (both Qiagen, Hilden, Germany). Extraction was performed according to the manufactureŕs instructions.

#### RIP (RNA immunoprecipitation)

HEK293 FT cells were cultured in DMEM (supplemented with 10% (v/v) FCS and 1% (v/v) Penicillin/Streptomycin) until they reached 70% confluency in a 10 cm cell culture dish. Transfection with the corresponding plasmids (RIG-I(WT) and RIG-I(I875A)) was conducted by using Lipofectamine2000. In short, 20 µg plasmid were mixed with 480 µl OptiMEM and 40 µl Lipofectamine2000 were mixed in a second tube with 460 µl OptiMEM. The plasmid mix was slowly added to the Lipofectamine2000 solution and briefly resuspended. After 10 minutes incubation at room temperature the solution was added dropwise to the cells in the 10 cm cell culture dish with 9 ml culture medium. The cells were incubated for 24h at 37°C with 5 % CO_2_ in an incubator. Subsequently, the cells were two times carefully washed with cold PBS and collected with a cell scraper. After centrifugation at 4°C for 5 minutes at 200 ×g the pellet was resuspended in 500 µl NP40 cell lysis buffer (50mM HEPES, pH 7.5, 150 mM KCl, 2 mM EDTA, 0.5 mM DTT, 0.5% (v/v) NP-40, protease inhibitors) and incubated on ice for 15 minutes. The solution was cleared by centrifugation at 4°C for 5 minutes at 20,000 ×g and the supernatant was transferred to a 2 ml reaction tube. To check for IP quality 50 µl were mixed with 500 µl TRIzol (Thermo Fisher Scientific) for input RNA and 50 µl were mixed with 10 µl 6xSDS loading dye for western blot analysis. The sample was filled up to 900 µl with NP40 cell lysis buffer for better binding during rotation. An amount of 30 µl anti-FLAG magnetic beads (Sigma M8823) per reaction were washed 2 times with 1 ml NP40 cell lysis buffer and resuspended in 100 µl NP40 cell lysis buffer. Afterwards, 100 µl anti-FLAG magnetic beads were added to each sample and incubated at 4°C on a rotating wheel for 16h. The anti-FLAG magnetic beads were washed 5 times with 1 ml NP40 cell lysis buffer and collected by a magnetic rack. 100 µl were saved for western blot IP analysis. The remaining anti-FLAG magnetic beads were collected, the supernatant was removed and 500 µl TRIzol was added. The sample was stored at -20°C. Subsequent RNA isolation and cDNA synthesis were conducted following the manufacturer’s instructions.

The data analysis was performed following the ΔCt method. In short, the IP values were normalized to GAPDH levels to obtain the ΔCt value. Afterwards, the fold enrichment was calculated by 2^-(ΔCt). Finally, the fold enrichments were normalized to the non-specific control (anti-magnetic FLAG beads in non-transfected WT cells) to determine the fold changes of all analyzed targets. Significances were calculated by student’s t test. The recovered yield of the RIP was calculated to check for IP quality. Western blot analyses using anti FLAG antibody (Sigma F1804) was performed to check for RIG-I recovery after IP.

#### Enzymatic treatment of RNA

FastAP Alkaline Phosphatase (Thermo Fisher Scientific) was used to remove 5’-phosphates from RNA samples to generate 5’OH-RNA. 0.25 µg RNA was incubated in 1 x FastAP reaction buffer containing 1 U FastAP for 30 min at 37°C.

RNA 5’Polyphosphatase (Lucigen, Middleton, USA) was used to generate 5’p-RNA from IVT 5’ppp-RNA. 5 µg RNA were incubated in 1 x polyphosphatase reaction buffer containing 20 U RNA 5’Polyphosphatase for 30 min at 37°C according to the manufacturer’s protocol.

1 µg of whole-cell RNA was treated with 1 µl RNase A (Thermo Fisher Scientific) in low salt (50 mM NaCl) or high salt (200 mM NaCl) for 30 min at 37°C.

#### qPCR

RNA from HEK Flp-In^™^ T-REx^™^ 293 and A549 cells was extracted using Zymo III columns (Zymo, Freiburg, Germany). Reverse transcription was performed using random hexamer primers (IDT) and RevertAid Reverse Transcriptase (Thermo Fisher Scientific). qPCRs (with the exception of SncmtRNA) was performed using 5x EvaGreen qPCR-Mix II (ROX) (Bio-Budget, Krefeld, Germany). Primers (Table S2) were validated with melting curve analysis and tested for efficiency using cDNA dilution. SncmtRNA qPCR was performed using Luna® Universal qPCR Master Mix (New England Biolabs). Primetime® Mini qPCR Assay FAM/ Iowa Black FQ labeled (Integrated DNA Technologies, Leuven, Belgium) (Table S2) was tested for efficiency using cDNA dilution. The SncmtRNA qPCR product, which spans a part of the loop connecting sense and antisense mt-rRNA was validated by sanger sequencing (Seqlab) to be 100% in sequence with DQ386868. qPCR was performed on a QuantStudio™5 Real-Time PCR System (Thermo Fisher Scientific) using standard Taqman Relative Quantification protocol.

#### Immunoblot

2 x 10^5^ HEK Flp-In^™^ T-REx^™^ 293 cells were seeded in 24-well plates and RIG-I expression was induced by addition of doxycycline. 72h later cells were lysed in 60 µl 1 x Laemmli-buffer (120 mM Tris pH 6.8, 4% (w/v) SDS, 20% (v/v) glycerol, 20 mM DTT and Orange G). Samples were sonificated and heated (95 °C, 5 min) prior to loading. Primary and secondary antibodies are listed in the KeyResourcestable. Blots were recorded on an Odyssey FC Dual-Mode Imaging system (LI-COR Biosciences, Bad Homburg, Germany).

#### Preparation of recombinant proteins

The coding sequence of RIG-I C-terminal domain (CTD) (802-925) and RIG-I (802-925, I875A) was cloned into pGEX-4T1 using *Nco*1 and *Eco*R1 restriction sites at the 5’ and 3’ end, respectively. Proteins were expressed in *E.coli* BL21(DE3) cells overnight at 22°C by induction with 0.4 mM IPTG. Cells were lysed by sonication using 20 mM Tris/HCl, pH 7.5, 200 mM NaCl, 2 mM MgCl_2_, 1 mM DTT, 12.5 µg/mL RNase A, spatula tip of lysozyme, and DNase. GST-RIG-I protein variants were affinity purified by GSTrap column (GE Healthcare). After sample loading, the column was washed with 20 mM Tris/HCl, pH 7.5, 1 M NaCl, 2 mM MgCl_2_, 1 mM DTT. Protein was eluted using 20 mM Tris/HCl, pH 7.5, 200 mM NaCl, 2 mM MgCl_2_, 1 mM DTT, and 10 mM GSH. For protein crystallization, pooled fractions were incubated with TEV protease (5 µg/mL) overnight at 4°C to cleave off the GST-tag. Sample was purified by S75 size exclusion column (GE Healthcare) and fractions containing RIG-I protein were concentrated to 10 mg/ml (RIG-I CTD(WT)) or 27 mg/ml (RIG-I CTD(I875A)), aliquoted in storage buffer, snap frozen in liquid nitrogen, and stored at −80°C.

#### Preparation of SncmtRNA in vitro transcripts

We hybridized two in vitro transcripts (IVT) to obtain a SncmtRNA like structure (see also Fig. 5A): Templates for in vitro transcription were generated by using 5’primers binding at position +10 (#1) or adding 3 nucleotides (GCC, #2) upstream of the SncmtRNA sequence (DQ386868) to obtain templates that are transcribable by T7 polymerases and that harbor a basepaired 5’end (like the original transcript) (see Fig. 5). The reverse primer binds at 809. For the second IVT template (#3) covering the loop and antisense part of SncmtRNA, a forward primer binding at 810 and reverse primer starting at 2374 were used. Purified PCR products were used for T7 in vitro transcription according to the manufacturers instructions (Thermoscientific). IVTs were treated with 5’polyphosphatase, producing 5’p-RNA from 5’ppp-RNA. IVT#1 or IVT#2 were hybridized with IVT#3.

#### Surface Plasmon Resonance Spectroscopy

Surface plasmon resonance experiments were performed using a Biacore 8K instrument (GE Healthcare). The flow system was cleaned using the maintenance “Desorb” function (Desorb Kit, GE Healthcare). The system was flushed with running buffer (20 mM Tris pH 7.4, 150 mM NaCl, 5 mM KCl, 1 mM CaCl_2_, 1 mM MgCl_2_, 0.05% Tween20) and all steps were performed at 25°C chip temperature. For GST-RIG-I CTD (wt or I875A) capturing, a CM5 sensor chip was covalently modified by an anti-GST polyclonal goat antibody (GST capture Kit, GE Healthcare). The surface of flow cell 1 and 2 was activated for 60 s with 50 mM NaOH (10 μL/min), followed by activation of both flow cells for 7 min with a 1:1 mixture of 0.1 M NHS (N-hydroxysuccinimide) and 0.1 M EDC (3-(N,N-dimethylamino) propyl-N-ethylcarbodiimide) (10 μL/min). Before amine-coupling, the flow system was washed with 1 M ethanolamine pH 8.0. The anti-GST antibody (30 μg/mL in 10 mM sodium acetate pH 5.0, 5 µL/min) was immobilized on flow cell 1 and 2. Surfaces were blocked with a 7 min injection of 1 M ethanolamine pH 8.0 (10 μL/min). For blocking of high affinity sites, recombinant GST was injected for three consecutive 180 s cycles (30 µL/min, 200 mM) followed by 120 s regeneration with 10 mM glycine pH 2.0 (30 µL/min). GST-RIG-I (wt or I875A) was immobilized for 240 s on flow cell 2 (4 µL/min, 2 µM) and recombinant GST for 90 s on reference flow cell 1 (10 µL/min, 200 nM). Before interaction measurements, the sensor chip was stabilized for 10 min with running buffer (30 µL/min).

For kinetic binding measurements the analyte was injected (30 μL/min, association: 60 s, dissociation: 240 s) over both flow cells at concentrations of 128, 256, 512, 1024, 2048 and 4096 nM (p/OH-dsRNA) or 0.25, 0.5, 1, 2, 4, 8 nM (3p-dsRNA). After each cycle, the surfaces were regenerated with a 120 s injection step of 10 mM glycine pH 2.1. Data were collected at a rate of 10 Hz. The data were double referenced by blank cycle and reference flow cell subtraction. Processed data were fitted by 1:1 interaction model using Biacore Insight Evaluation Software (version 2.0. 15.12933).

#### Crystallization, structure solution and refinement

The purified and concentrated RIG-I (wild type, 10.3 mg/ml) was stored in 20 mM Tris–HCl pH 7.5, 50 mM NaCl, 1 mM Dithiothreitol (DTT). Initial crystals of wild-type RIG-I in complex with RNA (OH-GFP2 Palindrome, 12mer) were obtained using the hanging drop vapor diffusion technique at 20°C by mixing 1 µl protein-complex (1:1 ratio) solution with 1 µl of the reservoir solution containing 0.1 M sodium cacodylate pH 6.5, 0.2 M NaCl, 22% Peg3350 at 10°C. After optimization, the best hexagonal-shaped crystals (dimensions of ∼0.3 x 0.3 x 0.4 mm) were obtained by using 100 mM Tris–HCl pH 8.0, 0.2M sodium chloride, 18% PEG 3350 as the reservoir and grew in ∼9-11 days.

The crystals of mutated RIG-I (I875A, 20mg/ml) in complex with RNA (p-GFP2 Palindrome, 12 mer) were obtained using 1:1 ratio of protein – RNA. The optimized crystals were grown using 0.2 M thiocyanate pH 8.4, 20% PEG 3350, 10% ethylene glycol and 10% MPD in the reservoir solution.

Diffraction data of wild type RIG-I in complex with RNA (OH-GFP2, 12mer) were collected at the Synchrotron SOLEIL Proxima-I, Saint-Aubin, France. Data were collected at 0.97857 Å wavelength and 100 K temperature. Diffraction data of mutant RIG-I(I875A) in complex with RNA (P-GFP2, 12mer) were collected at PX-II beamline at Swiss Light Source Villigen at 0. 9999097 Å wavelength and 100 K temperature using the EIGER detector.

The XDS package (Kabsch, 2010) (ver. 2019) was used to process, integrate and scale all the data. The structure of the 5’-p-dsRNA CTD I875A complex was solved by molecular replacement with PHASER (McCoy, as implemented in phenix v.1.17.1) using PDB-ID 3NCU as search model. The three monomers were found in three separate runs with a final TFZ-score of 29.3 and LLG of 1378.08. The structure was refined using phenix.refine (Liebschner et al., 2019, ver. 1.17.1) leading to R/Rfree values of 21.8/27.3. The 5’-OH-dsRNA CTD structure was also solved with PHASER (TFZ 28.4, LLG 926) and refined using phenix.refine with twin law (h,-h-k,-l; twin fraction 0.23) leading to R/Rfree values of 19.3/21.8. The geometry of the structures was checked and optimized with COOT (0.8) and MOLPROBITY (http://molprobity.biochem.duke.edu) (Emsley and Cowtan, 2004; Williams et al., 2018) (Suppl. Fig. 3A).

The XDS package (*Kabsch, 2010*) was used to process, integrate and scale all the data. The structure of the 5’-p-dsRNA CTD I875A complex was solved by molecular replacement with PHASER (McCoy) using PDB-ID 3NCU as search model. The three monomers were found in three separate runs with a final TFZ-score of 29.3 and LLG of 1378.08. The structure was refined using phenix.refine (Liebschner et al., 2019) leading to R/Rfree values of 21.8/27.3. The *5’-OH-dsRNA CTD* structure was also solved with PHASER (TFZ 28.4, LLG 926) and refined using phenix.refine with twin law (h,-h-k,-l; twin fraction 0.23) leading to R/Rfree values of 19.3/21.8. The geometry of the structures was checked and optimized with COOT and MOLPROBITY (Emsley and Cowtan, 2004; Williams et al., 2018) (Suppl. Fig. 3A).

### QUANTIFICATION AND STATISTICAL ANALYSIS

Statistical analysis was performed using GraphPad Prism 6 and 8 for Mac OS X. Statistical parameters are reported in the figure legends.

---

**Suppl. Figure 1:**
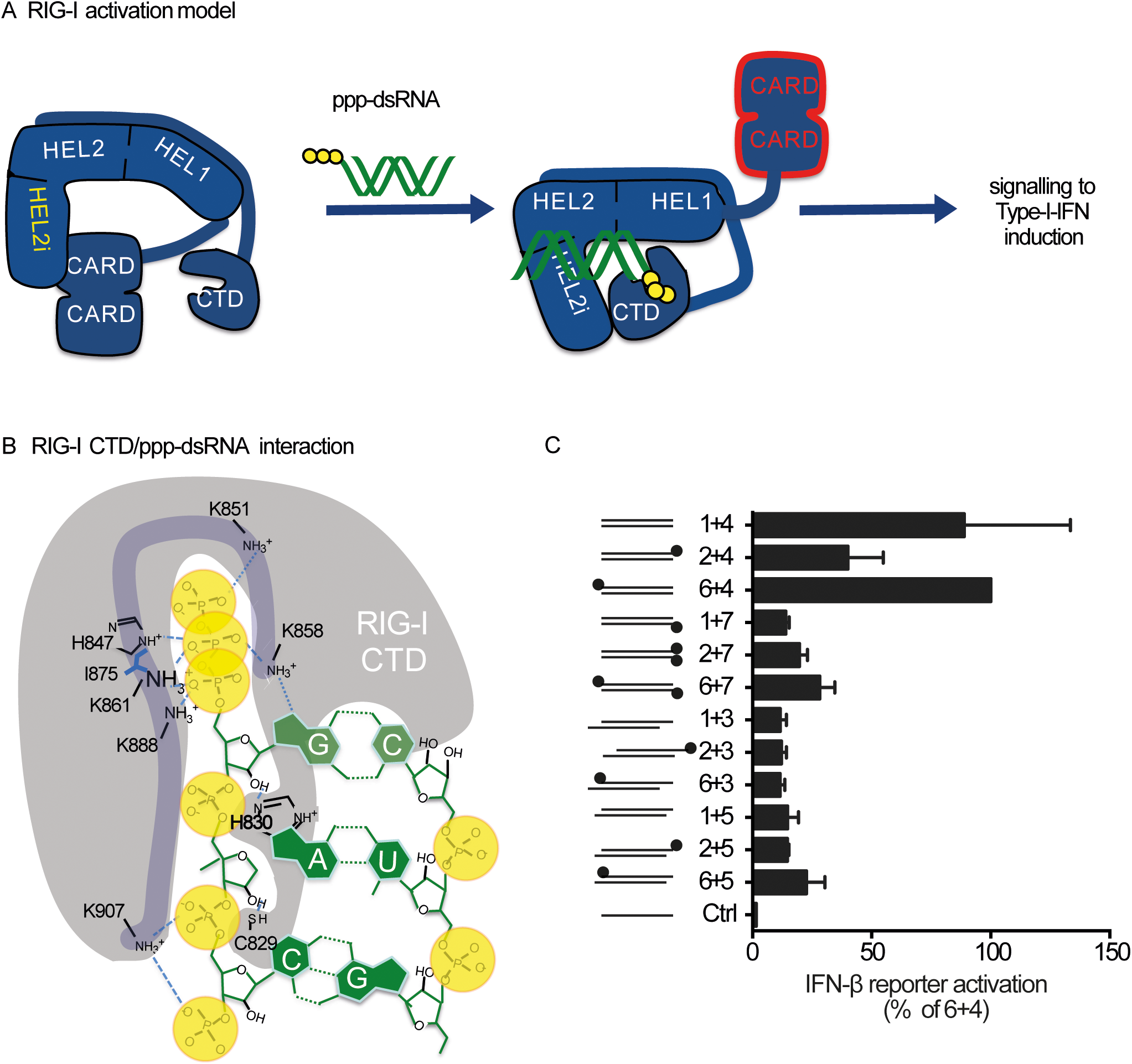
A) RIG-I activation model from Kowalinski et al., 2012: An inhibitory helicase subdomain (Hel-2i) binds the caspase activation and recruitment domains (CARDs) in this way suppressing CARD signaling. Upon RNA ligand binding by the C-terminal domain (CTD) the CARD domains are released to signal to type-I-IFN induction B) Scheme of amino-acid interaction with RNA in the RIG-I(WT) CTD C) HEK T-REx RM^-/-^ cells expressing RIG-I(WT) were stimulated with indicated synthetic RNA ligands (1 µg/ml). IFN-β promoter induction was determined by dual-luciferase assay 20 h after RNA transfection (n = 2, mean + SD).

**Suppl. Figure 2.**
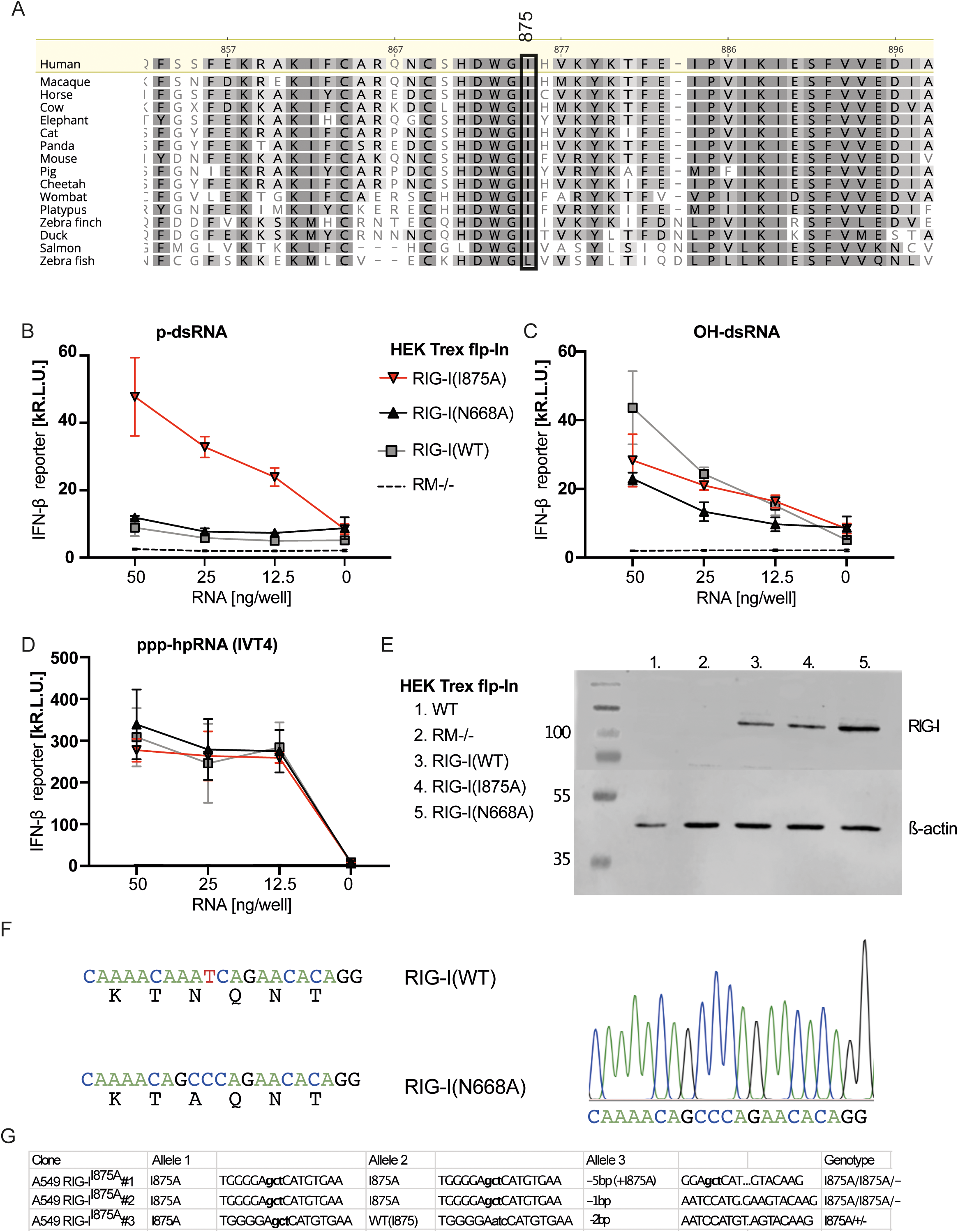
1875 is highly conserved among vertebrates. A) Amino acid alignment of the RIG-I sequences of 16 species. I875 is highlighted with a black box. Alignment was performed using Geneious Prime. B,C,D) HEK T-REx RM^-/-^ cells expressing RIG-I(WT), RIG-I(I875A) or RIG-I(N668A) upon addition of 0.1 ug/ml doxicycline were stimulated with p-dsRNA (B) =H-dsRNA (C) or ppp-hpRNA (D). IFN-β reporter activation was determined 20 h after stimulation (n=3). E) Expression of RIG-I was assessed by western blot. F) Sequence and construct sequencing results from the RIG-I(N668A) construct. G) Sequencing results from the endogenous DDX58 locus in A549 cells.

**Suppl. Figure 3:**
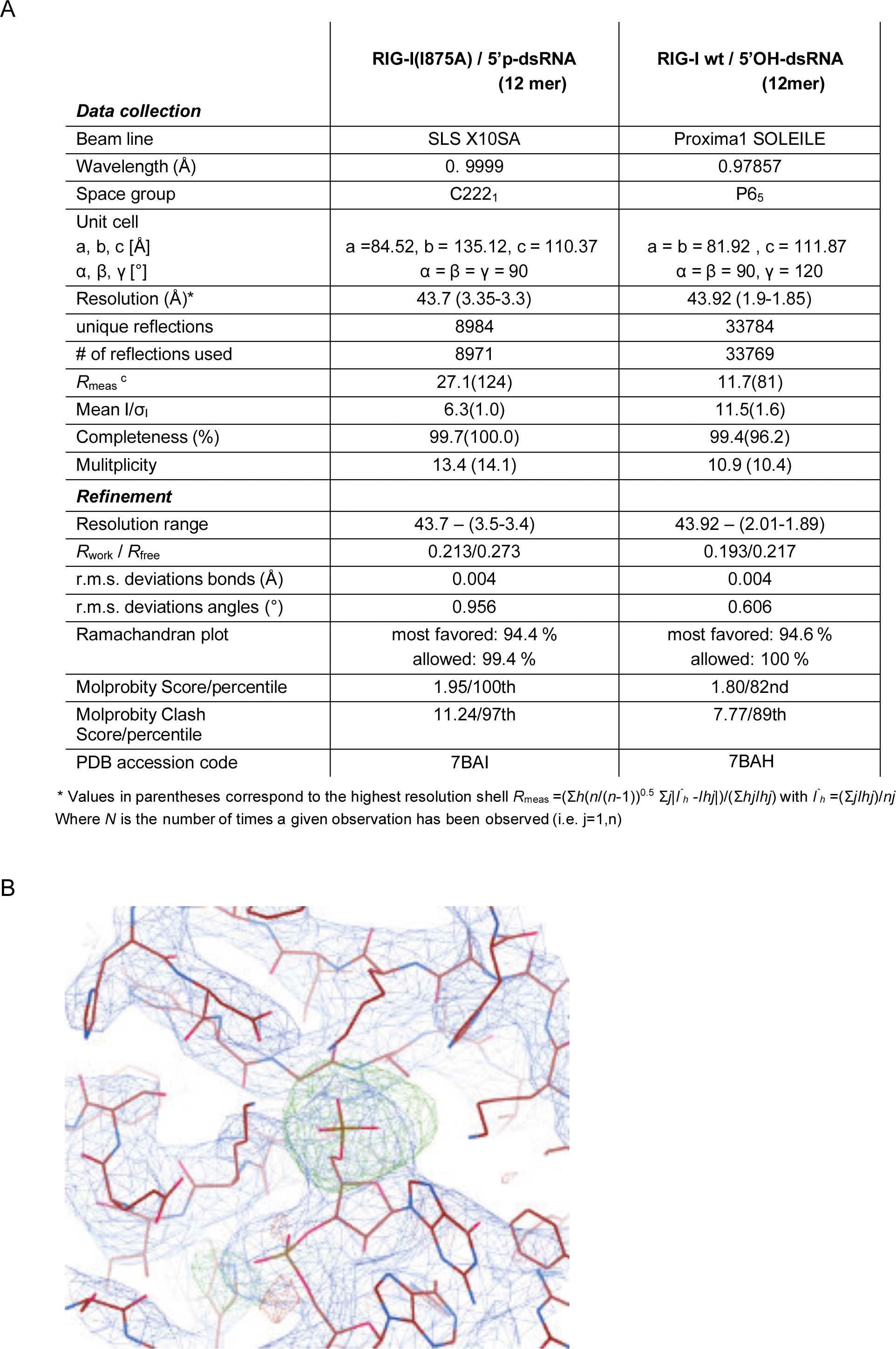
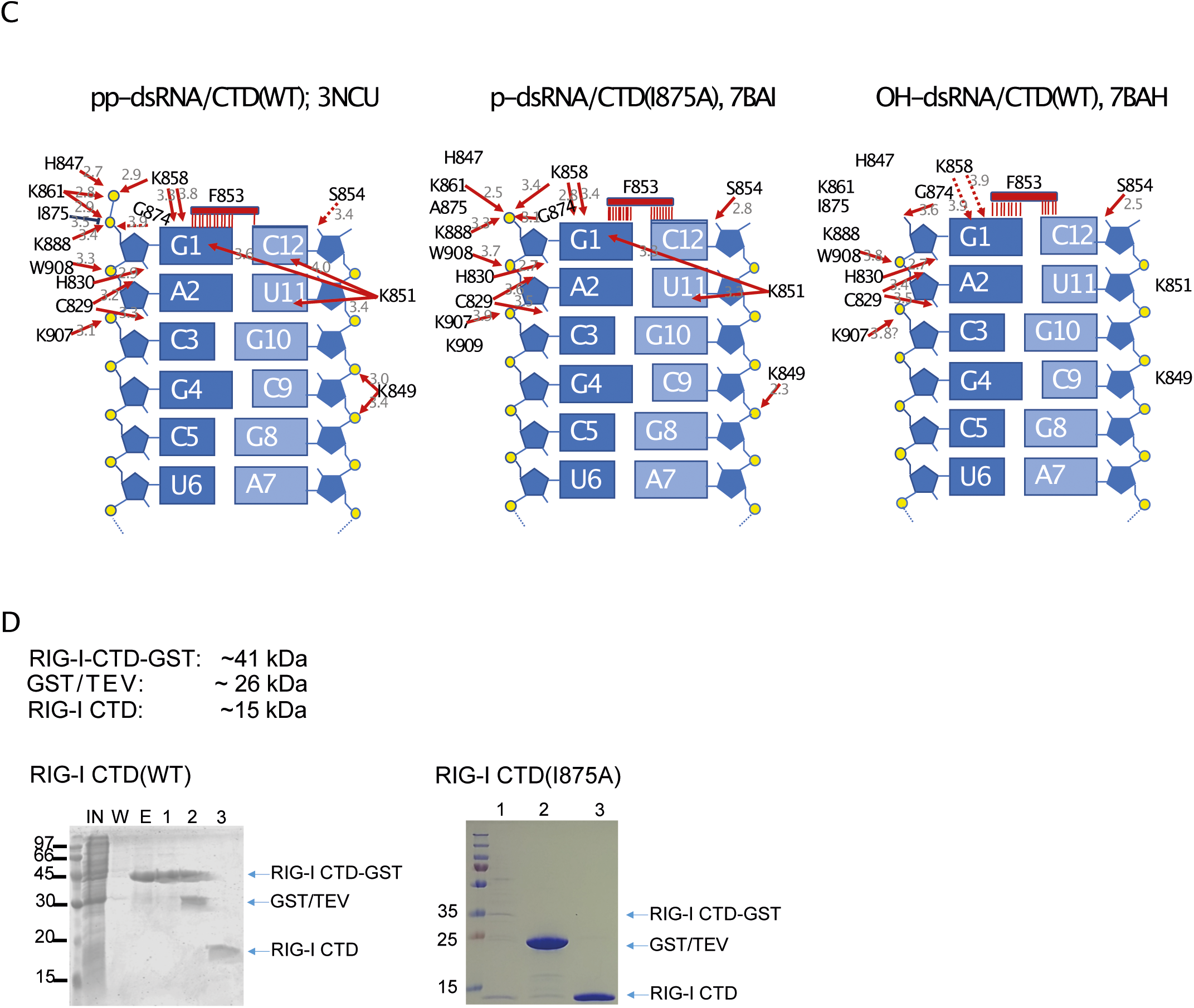
Crystal data. A) The structure of the 5’p-dsRNA CTD I875A complex was solved by molecular replacement with PHASER (McCoy, PHENIX) using PDB-ID 3NCU as search model. The three monomers were found in three separate runs with a final TFZ-score of 29.3 and LLG of 1378.08. The structure was refined using phenix.refine leading to R/Rfree values of 21.8/27.3. The 5’-OH-dsRNA CTD structure was also solved with PHASER (TFZ X.x, LLG Y.y) and refined using phenix.refine (Liebschner et al., 2019) with twin law (h,-h-k,-l; twin fraction 0.23) leading to R/Rfree values of 19.3/21.8. The geometry of the structures was checked and optimized with COOT and MOLPROBITY (Emsley and Cowtan, 2004; Williams et al., 2018). B) The omit map was calculated with phenix.refine using the simulated annealing option and by setting the occupancy of the phosphate group to zero. The blue mesh represents 2mFo-DFc density at 1.0 o-. The green and red meshes are mFo-DFc densities at 3.0 o-. The S’p-dsRNA CTD 187SA structure is shown as a brown stick model. C) Scheme of amino acid-RNA interactions within the indicated structures. Binding distances are depicted in angstrom. D) Comassie blue stained PAGE of recombinant RIG-I CTD-GST, cleaved or uncleaved behind the GST by TEV (WT, left; I875A, right).

**Suppl. Figure 4:**
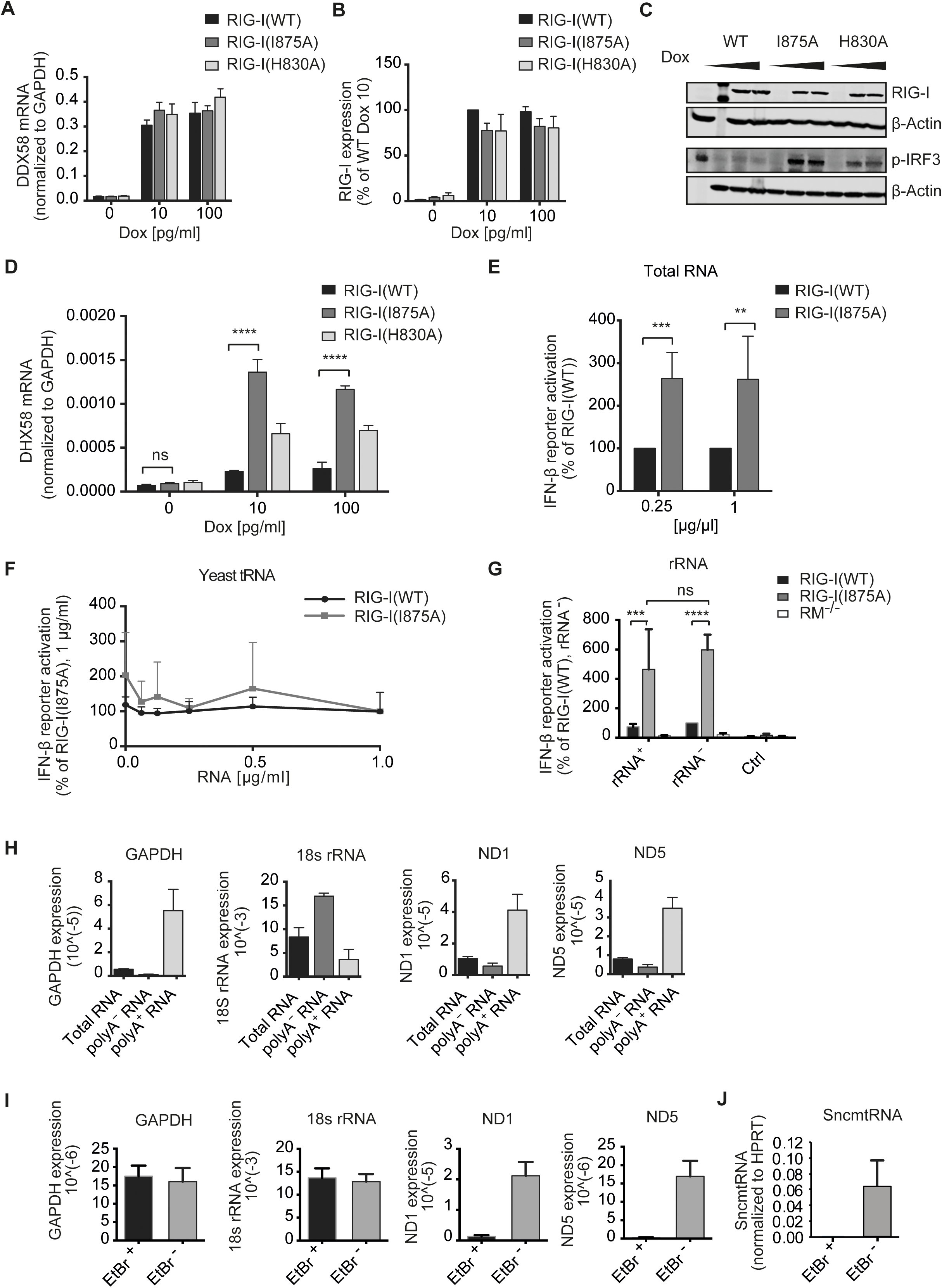
Identification of endogenous RIG-I(I875A) stimulating RNA species. A-D) RIG-I(WT), RIG-I(I875A) or RIG-I(H830A) expression was induced with indicated Doxycycline concentrations (Dox) in HEK T-REx RM^-/-^ cells in the absence of exogenous RIG-I stimuli. A,D) RNA was extracted 72h later, and mRNA expression of DDX58 (RIG-I) (A, n=3-4) and DHX58 (D, n=3) were analyzed by qPCR and normalized to GAPDH (mean + SEM, D, two-way ANOVA, Tukey’s post test). B) Protein was extracted 72 h later and RIG-I expression was analyzed by western blot and normalized to β-Actin (n = 4, mean + SEM). C) One representative western blot experiment from B) / Fig. 4 B) is shown. E) Total RNA was extracted from A549 cells using TRIzol. HEK T-REx RM^-/-^ cells expressing RIG-I(WT) or RIG-I(I875A) were stimulated with extracted total RNA (0.25 or 1 µg/ml). IFN-β reporter activation was determined 20h after RNA transfection and is depicted as percentage of the signal of RIG-I(WT) (n = 8 (0.25 µg/ml), n = 9 (1 µg/ml), mean + SEM, paired t-test). F) HEK T-REx RM^-/-^ cells expressing either RIG-I(WT), RIG-I(I875A) or RIG-I(H830A) were stimulated with indicated concentrations of yeast tRNA. IFN-β reporter activation was determined 20h after RNA transfection and is depicted as the percentage of the signal of RIG-I(I875A), 1 µg/ml tRNA (n = 2, mean + SD). G) Total RNA from A549 cells was fractionated into a rRNA enriched (rRNA^+^) and rRNA depleted (rRNA^-^) fraction. Fractionated RNA was transfected into HEK T-REx RM^-/-^ cells expressing RIG-I(WT) or RIG-I(I875A) (1 µg/ml). IFN-β reporter activation was determined 20h after RNA transfection and is depicted as percentage of the signal of RIG-I(WT) stimulated by rRNA^-^ (n = 4, mean + SD, two-way ANOVA, Sidak’s post-test). H) Total RNA, polyA^+^ and polyA^-^ RNA from A549 cells were analyzed via qPCR for content of indicated RNAs (n = 5, mean + SEM). Expression is depicted as 2^(-Ct)^. I) RNA EtBr^+^ and RNA EtBr^-^ was analyzed via qPCR for content of indicated RNAs (n=8-11, mean + SEM). Expression is depicted as 2^(-Ct)^. J) RNA EtBr^+^ and RNA EtBr^-^ from A549 cells was analyzed via qPCR for content of SncmtRNA (n=12, mean + SEM). Stastical significance is indicated as follows: *** (p < 0.001), **** (p < 0.0001), ns: not significant. Dox: Doxycycline

## KEY RESOURCES TABLE

| REAGENT or RESOURCE | SOURCE | IDENTIFIER |
| --- | --- | --- |
| **Antibodies** |  |  |
| Rabbit monoclonal anti-RIG-I (D14G6) | Cell Signaling Technology | Cat# 3743S;<br>RRID:AB_2269233 |
| Rabbit monoclonal anti-phospho-IRF-3 (Ser396) (4D4G) | Cell Signaling Technology | Cat# 4947S;<br>RRID:AB_823547 |
| Mouse monoclonal anti-β-actin | LI-COR Biosciences | Cat# 926-42212;<br>RRID:AB_2756372 |
| IRDye 800CW Goat anti-rabbit | LI-COR Biosciences | Cat# 926-32211;<br>RRID:AB_621843 |
| IRDye 680RD Goat anti-mouse | LI-COR Biosciences | Cat# 926-68070;<br>RRID:AB_10956588 |
| **Bacterial and Virus Strains** |  |  |
| *E. coli* |  | BL21(DE3) |
| *E. coli* | Thermo Fisher Scientific | Stbl3 |
| **Chemicals, Peptides, and Recombinant Proteins** |  |  |
| Ethidium bromide solution | Merck | Cat# E1510 |
| Uridine | Merck | Cat# U3003 |
| Sodium pyruvate | Thermo Fisher Scientific | Cat# 11360070 |
| MEM Non-Essential Amino Acids Solution | Thermo Fisher Scientific | Cat# 11140050 |
| tYeast-RNA | Thermo Fisher Scientific | Cat# AM7119 |
| Lipofectamine 2000 Reagent | Thermo Fisher Scientific | Cat# 11668019 |
| NTC buffer | Macherey Nagel | Cat# 740654.100 |
| FastAP Alkaline Phosphatase (Thermo Fisher Scientific) | Thermo Fisher Scientific | Cat# EF0651 |
| RNA 5' Polyphosphatase (Lucigen) | Lucigen | Cat# RP8092H |
| RNase A (Thermo Fisher Scientific) | Thermo Fisher Scientific | Cat# EN0531 |
| TRIzol™ | Thermo Fisher Scientific | Cat# 15596018 |
| **Critical Commercial Assays** |  |  |
| Ribominus™ Human/Mouse Transcriptome Isolation Kit | Thermo Fisher Scientific | Cat# K155002 |
| NucleoSpin® Gel and PCR Clean-Up Kit | Macherey Nagel | Cat# 740609.50 |
| TranscriptAid™ T7 High Yield Transcription Kit | Thermo Fisher Scientific | Cat# K0441 |
| IFN alpha Human ELISA Kit | Thermo Fisher Scientific | Cat# BMS216 |
| miRNeasy Mini Kit | Qiagen | Cat# 217004 |
| RNeasy MiniElute Cleanup Kit | Qiagen | Cat# 74204 |
| NucleoTrap® mRNA Kit (Macherey Nagel) | Macherey Nagel | Cat# 740655 |
| **Deposited Data** |  |  |
| RIG-I CTD (I875A) – p-dsRNA structure | Protein Data Bank | 7BAI |
| RIG-I CTD WT – OH-dsRNA structure | Protein Data Bank | 7BAH |
| **Experimental Models: Cell Lines** |  |  |
| A549 cells |  |  |
| Flp-In™ T-Rex™ 293 Cell Line | Thermo Fisher Scientific | Cat# R78007 |
| Flp-In™ T-Rex™ 293 RIG-I/MDA5 -/- Cell Line | This paper | N/A |
| Flp-In™ T-Rex™ 293 RIG-I/MDA5 -/- Cell Line + RIG-I WT | This paper | N/A |
| Flp-In™ T-Rex™ 293 RIG-I/MDA5 -/- Cell Line + RIG-I (I875A) | This paper | N/A |
| Flp-In™ T-Rex™ 293 RIG-I/MDA5 -/- Cell Line + RIG-I (H830A) | This paper | N/A |
| Oligonucleotides |  |  |
| synthetic RNA oligonucleotides | Biomers.net, see Table S1 for sequences | see Table S1 |
| synthetic DNA oligonucleotides | Integrated DNA Technologies, see Table S2/S3 for sequences | see Table S2/S3 |
| Recombinant DNA |  |  |
| pOG44 Flp-Recombinase Expression Vector | Thermo Fisher Scientific | Cat# V600520 |
| PcDNA™5/FRT Mammalian Expression Vector | Thermo Fisher Scientific | Cat# V601020 |
| PcDNA5/FRT-RIG-I FL WT | This Paper | N/A |
| PcDNA5/FRT-RIG-I FL I875A | This Paper | N/A |
| PcDNA5/FRT-RIG-I FL H830A | This Paper | N/A |
| CAS9-gRNA expression plasmid | Thomas Zillinger | N/A |
| pMAX-CRISPR (Cas9-2A-EGFP, U6-sgRNA expression plasmid) | Thomas Zillinger | N/A |
| pIFN-β-G(aussia)-Luc(iferase) | Eicke Latz Lab | N/A |
| pLenti-Ef1α-F(irefly)-Luc(iferase) | Thomas Zillinger | N/A |
| Software and Algorithms |  |  |
| PRISM 6 | GraphPad Software | RRID:SCR_002798 |
| Geneious Prime | Geneious biologics | RRID:SCR_010519 |
| ImageStudioLite | LI-COR Biosciences | RRID:SCR_013715 |
| PHASER | McCoy, PHENIX |  |
| COOT, MOLPROBITY |  |  |
| phenix.refine | PHENIX |  |

---

**Table S1:**
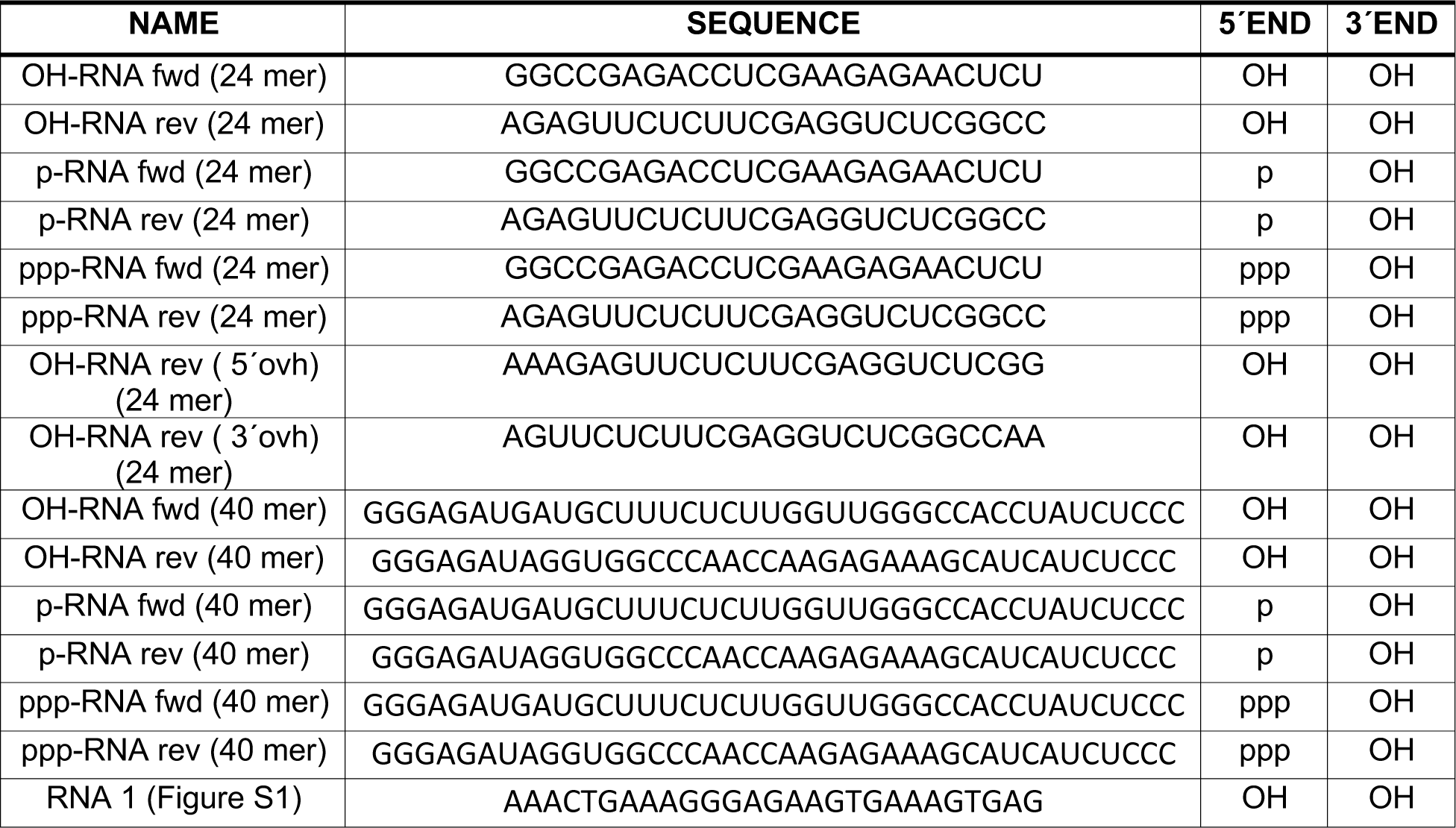
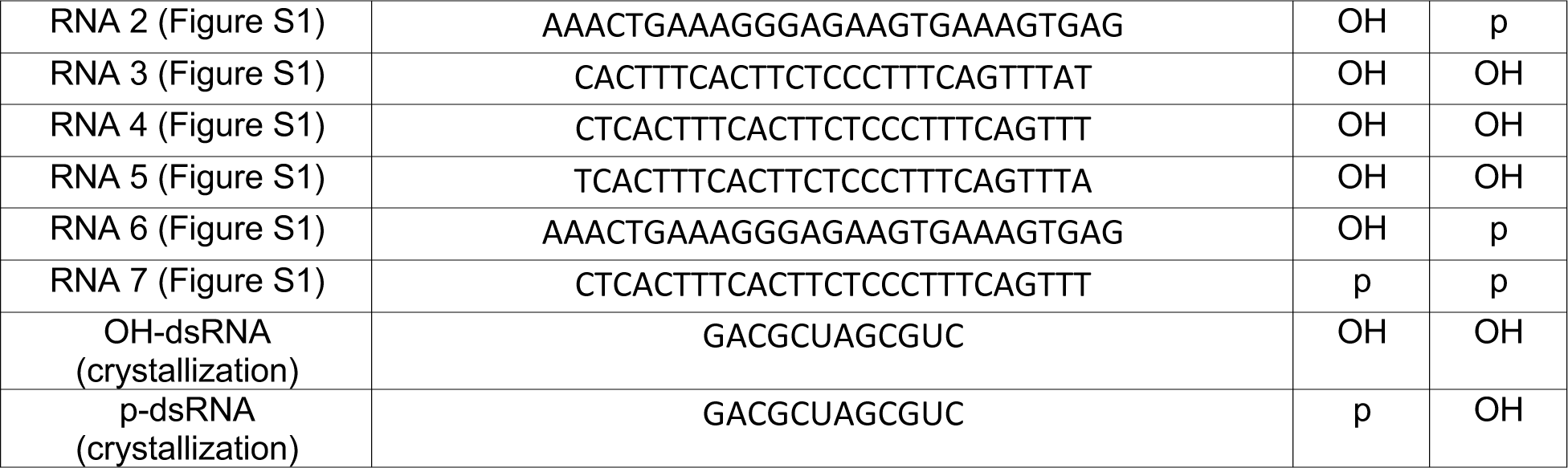
RNA oligonucleotide sequences.

**Table S2:**
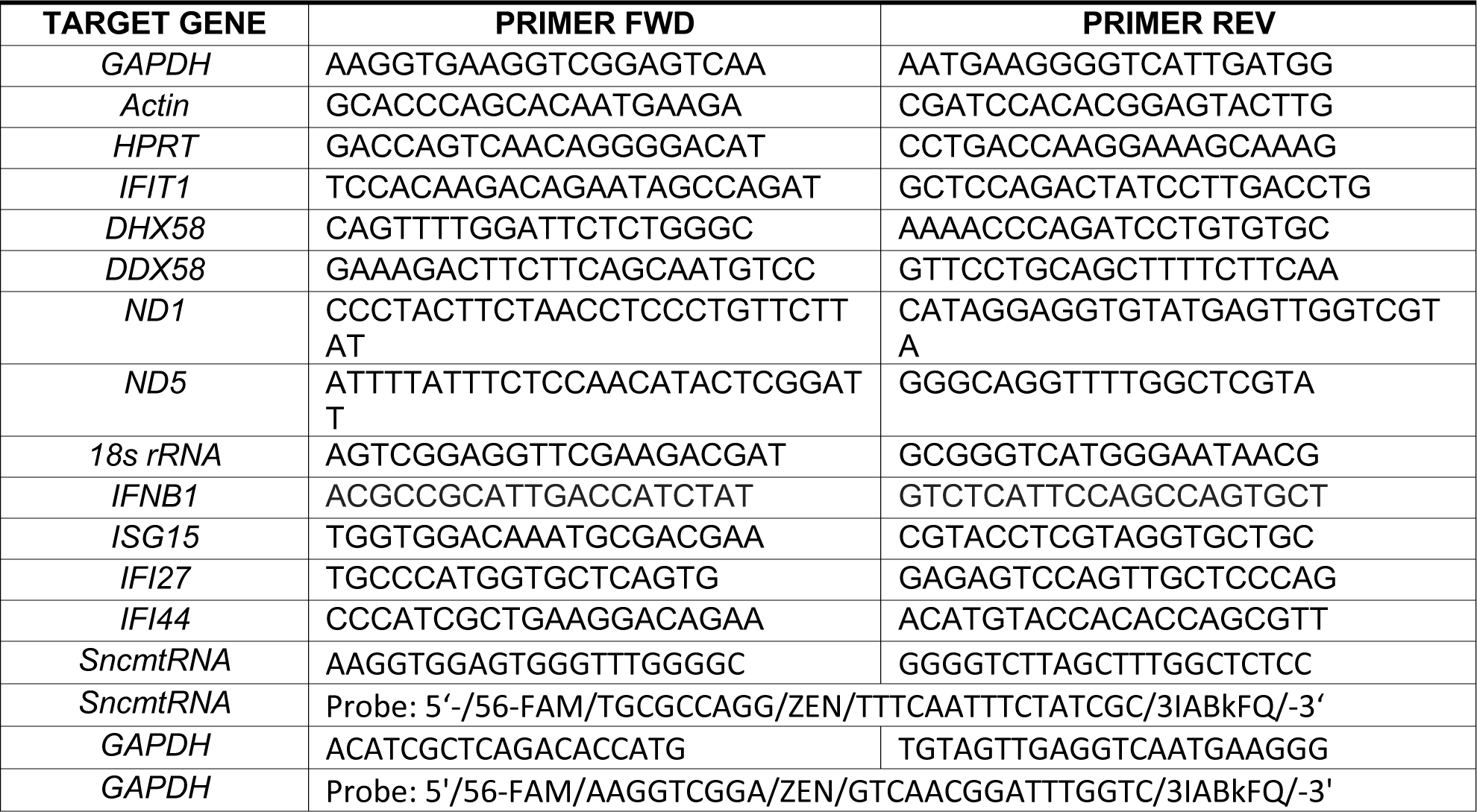
qPCR primer sequences.

**Table S3:**
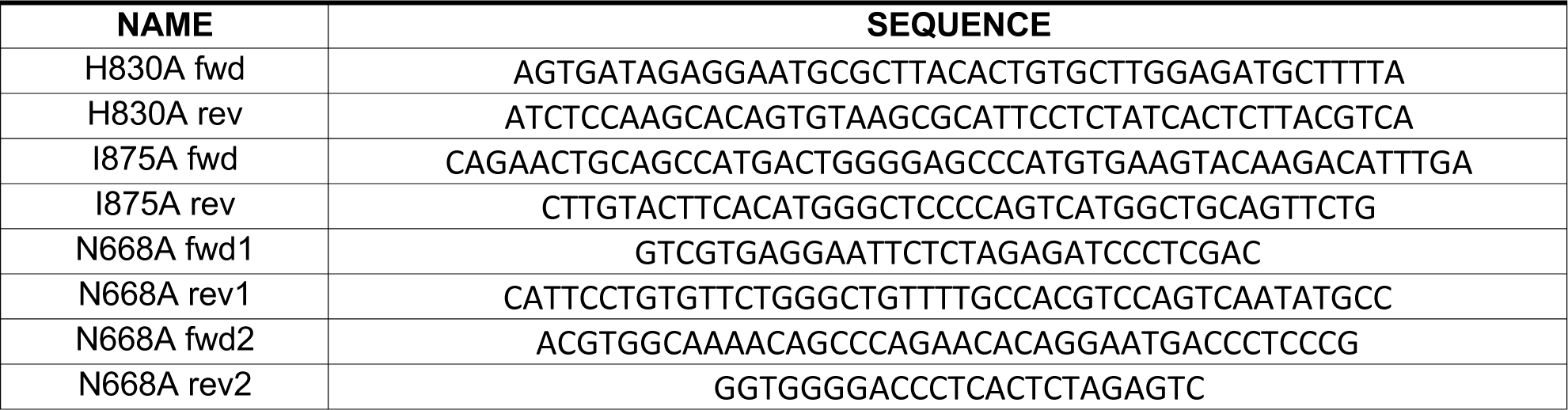
Cloning primer sequences.

**Table S4:**
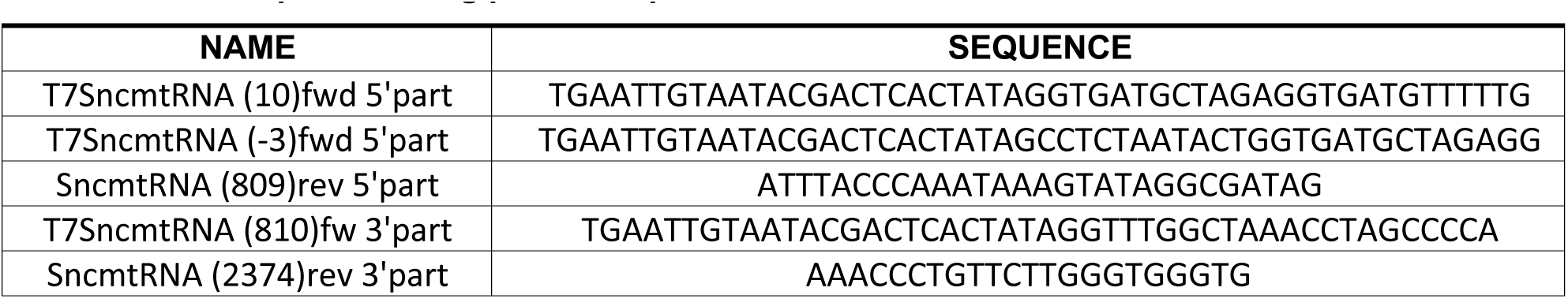
IVT template cloning primer sequences.

